# Neuroinflammation in the normal appearing white matter of multiple sclerosis brain causes abnormalities at the node of Ranvier

**DOI:** 10.1101/2020.06.10.142281

**Authors:** Patricia Gallego Delgado, Rachel James, Eleanor Browne, Joanna Meng, Swetha Umashankar, Carmen Picon, Nicholas D. Mazarakis, A. Aldo Faisal, Owain W. Howell, Richard Reynolds

## Abstract

Changes to the structure of nodes of Ranvier in the normal-appearing white matter (NAWM) of MS brains are associated with chronic inflammation. We show that the paranodal domains in MS NAWM are longer on average than control, with Kv1.2 channels dislocated into the paranode. These pathological features are reproduced in a model of chronic meningeal inflammation generated by the injection of lentiviral vectors for the lymphotoxin-α (LTα) and interferon-γ (IFNγ) genes. We show that tumour necrosis factor (TNF), IFNγ and glutamate can provoke paranodal elongation in cerebellar slice cultures, which could be reversed by an NMDA blocker. When these changes were inserted into a computational model to simulate axonal conduction, a rapid decrease in velocity was observed, reaching conduction failure in small diameter axons. We suggest that glial cells activated by proinflammatory cytokines can produce high levels of glutamate, which triggers paranodal pathology, contributing to axonal damage and conduction deficits.

## Introduction

Multiple Sclerosis (MS) is a neuroinflammatory disease of the central nervous system (CNS) characterised by focal and diffuse areas of inflammation, axonal degeneration and loss, demyelination and gliosis (Friese et al., 2014). Although the focus of MS research has for a long time been on the demyelinating lesions, neuronal damage and axonal loss are now recognised as early and persistent factors in MS pathology (Trapp et al, 1998) and may be the best predictors of long-term neurological decline (Reynolds et al., 2011). It has been suggested that activated microglia and macrophages surrounding myelinated stressed axons (Trapp et al., 1998) could initiate swelling around the nodes of Ranvier, leading to mitochondrial pathology and subsequent focal axonal degeneration (Nikic et al., 2011). The lesion free normal-appearing white matter (NAWM) in progressive MS is actually highly abnormal and contains chronically activated microglia, degenerating axons, dysfunctional astrocytes and a compromised blood-brain barrier (BBB) (Allen et al., 2001; Kirk et al., 2003; Zeiss et al., 2007; Dutta et al., 2018). Additionally, MRI studies have shown abnormalities in NAWM regions, especially in chronic progressive patients with long disease duration (Miller et al., 2002; Evangelou et al., 2004).

Axonal degeneration in the NAWM could be exacerbated by structural alterations at the nodes since they are critical elements in maintaining fast and efficient saltatory action potential (AP) conduction. The interaction between the Caspr1, contactin and neurofascin 155 (Nf155) proteins, and the cytoskeletal proteins, ankyrinB, αII and βII spectrins, contribute to the formation of the paranodal junctions (PNJ) (Peles et al., 1997; Tait et al., 2000; Simons et al., 2014; Arancibia-Carcamo and Attwell, 2014). The intricate and tight molecular interactions between the oligodendrocyte and the axon at these myelin free points are vital for restricting movement of membrane proteins between the various nodal zones and reducing the flow of current under the myelin sheath, but also make the PNJ particularly vulnerable to immune-mediated pathological alterations. Post-mortem tissue studies of MS NAWM have shown an increase in the length of Nf155 stained paranodal structures, and a partial dislocation of juxtaparanode Kv1 channels towards the node in a proportion of axo-glial junctions (Howell et al., 2010). However, the functional significance and mechanisms underlying the paranodal/nodal disorganisation in MS NAWM are unclear. Live laser-scanning coherent anti-Stokes Raman scattering (CARS) imaging of spinal cord myelin of rodent axon tracts exposed to elevated glutamate levels, pathological Ca^2+^ influx and calpain1 activation, has demonstrated paranodal splitting (Fu et al., 2009, 2011; Huff et al., 2011). Animal studies performed with conditional knockouts of the paranodal proteins Caspr1 (Bhat et al., 2001), Nf155 (Zonta et al., 2008), βII spectrin (Zhang et al., 2013) and 4.1.B protein (Cifuentes-Diaz et al., 2011; Buttermore et al., 2011) showed a lack of tight septate junctions, an increased peri-axonal space, dislocation of the juxtaparanodal voltage-gated channels Kv1 towards the PNJ, and functional alterations such as motor tremors and reduced conduction velocities. Taken together, this suggests a possible model of molecular paranodal disorganisation due to Ca^2+^ accumulation mediated by glutamate activation of the NMDA receptors located in the cytoplasmic processes of the oligodendrocyte at the PNJ (Salter and Fern, 2005; Karadottir et al., 2005; Micu et al., 2016). Multiple magnetic resonance spectroscopy (MRS) studies have demonstrated elevated glutamate levels in both acute MS lesions and NAWM tissue (Srinivasan et al., 2005; Tisell et al., 2013; Azevedo et al., 2014). Pro-inflammatory cytokines, such as TNF, can stimulate microglia in an autocrine/paracrine manner to induce glutamate release (Takeuchi et al., 2006; Yawata et al., 2008), as well as blocking astrocytic glutamate transporters crucial for glutamate homeostasis (Chao and Hu, 1994; Wang et al., 2003; Sitcheran et al., 2005). Recent data from human tissue studies has shown the presence of increased levels of TNF signaling in the MS brain (Magliozzi et al., 2019).

Here, we report that disruptions of the Caspr1 expressing PNJ structures and K_v_1 displacement are present in the NAWM of MS brains at regions remote from lesions and are accompanied by axonal changes, which could be reproduced in a rat model of chronic meningeal inflammation that leads to chronic microglial activation throughout the brain. Paranodal disruption correlated with the presence of activated microglia, suggesting a mechanism of axonal injury that starts at the paranode independently of the demyelination process. Furthermore, we demonstrate that similar PNJ pathology could be induced in ex-vivo cerebellar slice cultures by the activation of microglia with IFNγ and TNF or LTα and resultant glutamate release. Finally, we used biophysical simulations to systematically explore the effects on conduction of a range of paranodal and juxtaparanodal structural alterations observed in the human tissue and animal model.

## Results

### The PNJ structure is disrupted in post-mortem MS NAWM

In order to characterize PNJ pathology present in human MS NAWM tissue and its relationship to local microglial activation and axonal cytoskeleton disruption, NAWM regions of interest at least 4-5mm away from a demyelinating lesion were carefully selected from snap-frozen tissue blocks incorporating the cerebral peduncle and the precentral gyrus, both of which have a high density of longitudinal axons. Immunofluorescence with antibodies to the myelin protein, myelin oligodendrocyte glycoprotein (MOG), confirmed the myelin integrity (figure 1A) and HLA-DR (MHC class II) antibodies confirmed the presence of microglia with an activated morphology (thicker and shorter processes) (figure 1B). Immunofluorescence for Caspr1 localised at the paranodes demonstrated that Caspr1-stained paranodes (2.8 ± 1.1 μm) were significantly 21.7% longer on average in the MS NAWM tissue than in the non-neurological control tissue (2.3± 0.8 μm, figure 1C-G). Furthermore, 43% of the paranodes in the MS tissue were longer than the 75% percentile of the control group (2.7μm) and 11.14% were longer than 4μm, compared to the 4.1% in the non-neurological controls (figure 1G). There was little difference between the two brain areas examined; the paranodal lengths in the cerebral peduncles of the mesencephalon were 2.81±0.019μm long and 12% of them were >4μm, compared to those in the precentral gyrus which were 2.79± 0.017μm long and 10% of them were >4μm (Supplementary Figure1). This suggests that the length of the paranodes, defined by Caspr1 labelling, was disrupted in a significant proportion of axons in MS brains compared to non-neurological controls.

**Figure 1:**
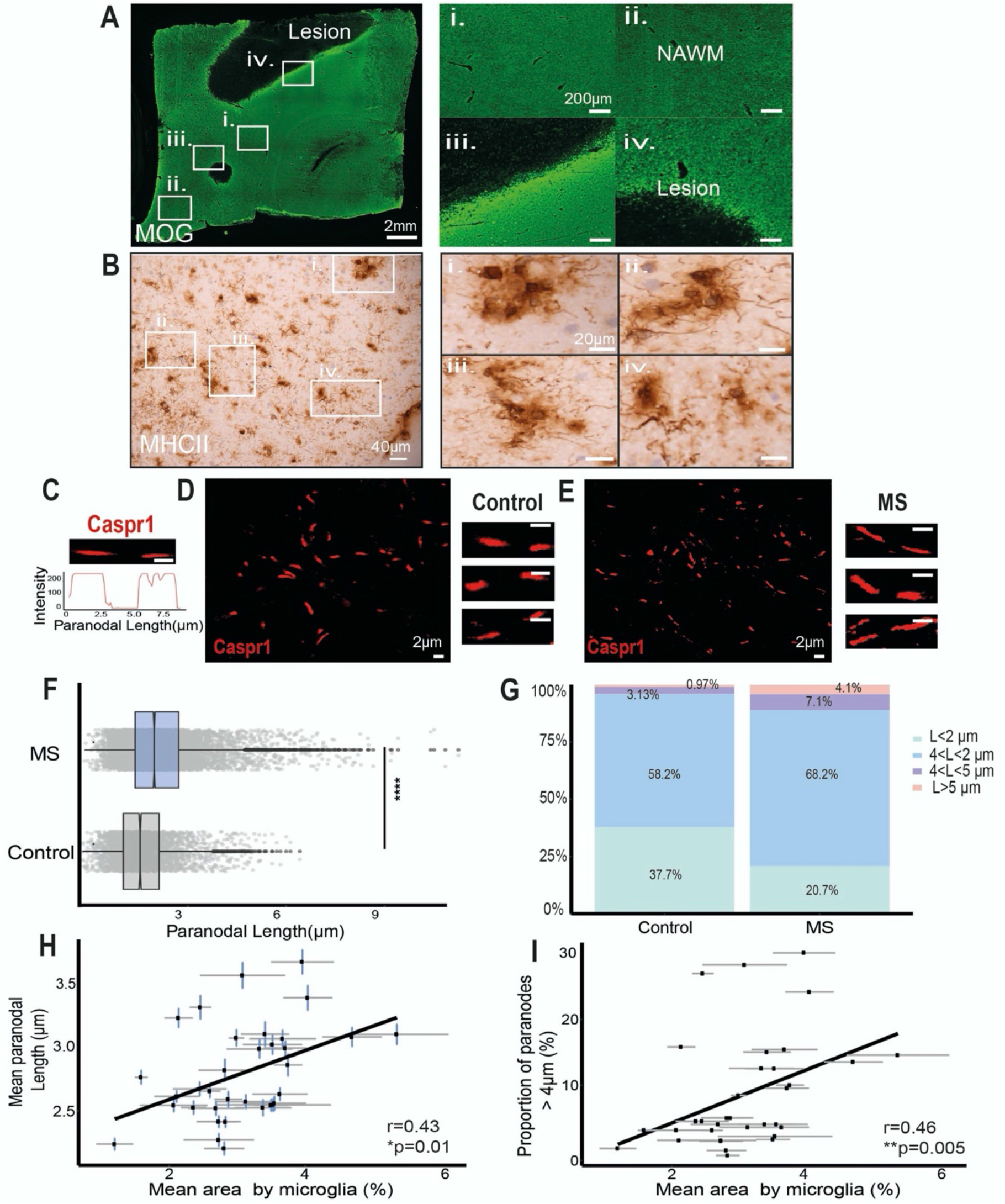
MS NAWM regions contained a larger proportion of elongated paranodes associated with activated microglia. (A) Anti-MOG-myelin immunofluorescence image. NAWM regions of interest (ROIs) were selected (i,ii) localised away from demyelinating lesions (iii, iv). (B) Process bearing anti-HLA-DR+ microglia with an activated morphology were found throughout the NAWM ROIs (i,ii).(C) Confocal image of single Caspr1-stained paranode in cross-section and its intensity profile. (D, E) Confocal images of paranodes from human post-mortem non-neurological control (D) and NAWM MS tissue (E). (F) Significantly different distributions of Caspr1+ paranodal lengths in NAWM MS tissue in comparison to non-neurological control tissue (p<0.0001, Mann-Whitney test). (G) NAWM MS tissue contained a larger proportion of Caspr1+ paranodes longer than 4 μm and 5μm than the control tissue. (H) The mean paranodal length per block correlated with the mean area occupied by HLA-DR+ microglia/ macrophages (r=0.43, *p<0.5, Spearman’s rank correlation test). (I) The mean area occupied by HLA-DR+ microglia/macrophages per block correlated with the proportion of paranodes longer than 4 μm (r=0.46, **p<0.01, Spearman’s rank correlation test).

### PNJ disruption is associated with microglial activation and axon stress

Chronic activation of microglia and axonal degeneration are two of the main pathological features of NAWM tissue in progressive MS. Therefore, we examined their relationship to paranodal length as a marker of paranodal axo-glial disruption. The mean area occupied by HLA-DR+ labelled microglia was obtained as a measurement of microglial activation. Moderate significant correlations were found between mean paranodal length and the proportion of paranodes >4μm and the mean microglial area (r=0.46 and r=0.43, *p<0.05, figure 1H,I). Double immunofluorescence labelling with the SMI32 antibody, which labels de-phosphorylated neurofilament proteins, an indicator of axon stress, and Caspr1 to indicate paranodal length (figure 2A), demonstrated that SMI32+ axons had longer Caspr1-stained paranodes on average (mean = 3.89 ± 0.1μm) than SMI32- (mean = 2.49 ± 0.07μm, figure 2B) and non-neurological controls (mean = 2.33 ± 0.01μm). Furthermore, the average paranodal length for SMI32+ axons was 66.9% longer than the control paranodal length and 56.2% longer than SMI32-axon paranodal length (figure 2B), indicating a strong relationship between the altered paranodes and axonal stress.

**Figure 2:**
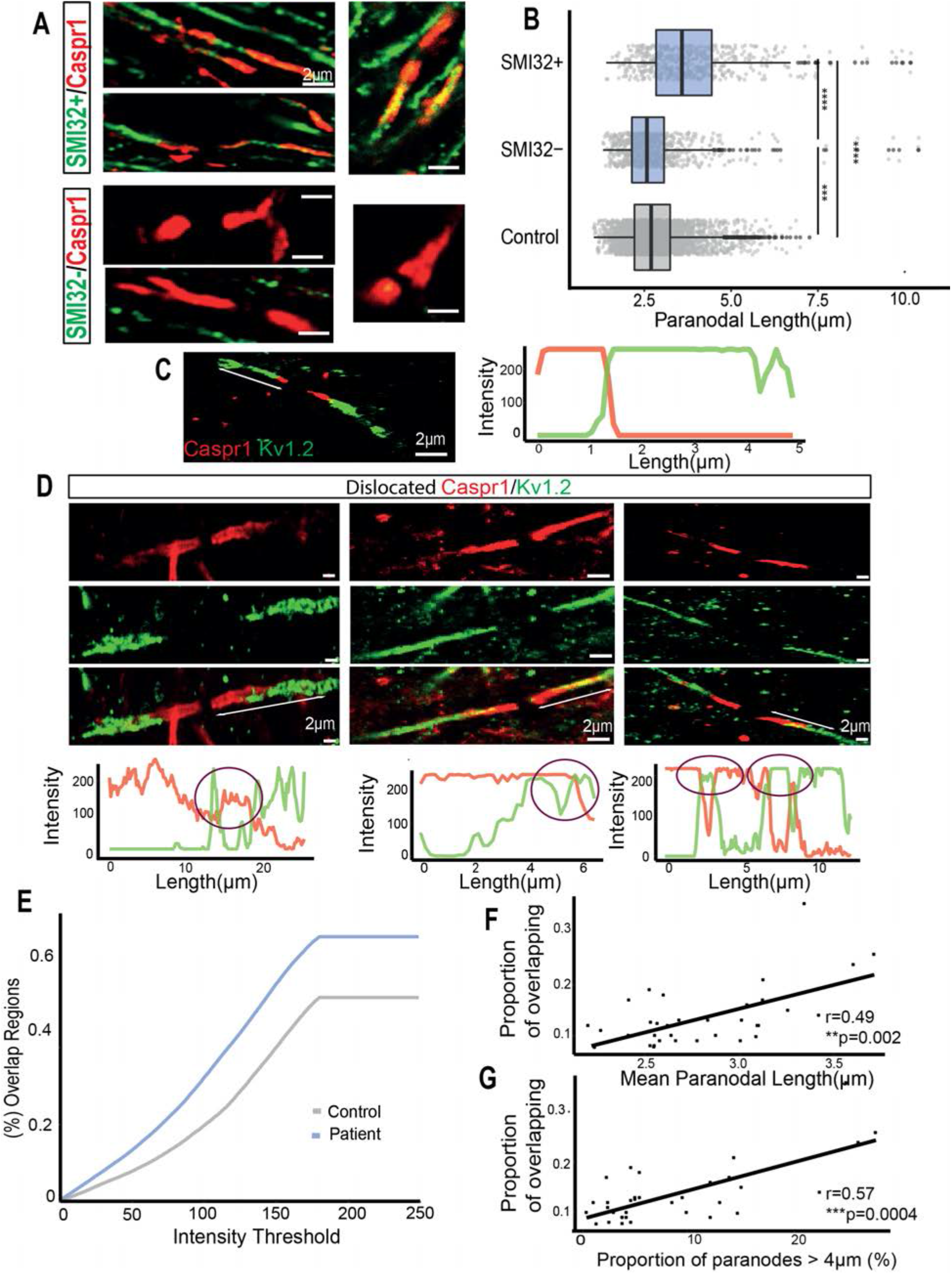
Paranodal elongation has been associated with SMI32+ axons and the dislocation of juxtaparanodal voltage-gated K_v_1.2 channels. (A) Confocal images of long Caspr1+ paranodes co-stained with SMI32 antibody. SMI32+ axons characterized by dephosphorylated neurofilaments had elongated paranodes. (B) Paranodal length distributions of SMI32+ and SMI32-axons from MS NAWM and non-neurological control tissue (**** p < 0.0001, Mann-Whitney test). (C) Confocal image from a node showing the expression of Caspr1 in the paranode (red) and K_v_1.2 channels in the juxtaparanodes (green) do not overlap under non-pathological conditions, and the respective RGB profile. (D) Confocal images from nodes where Caspr1 and K_v_1.2 are colocalising, and therefore possibly being affected by MS neuropathological conditions, and their intensity RGB profiles. The purple circles denote regions where both signals are colocalising: overlapping regions. (E) When the difference between Caspr1 and K_v_1.2 signals was smaller than a variable intensity threshold, we considered that point as an overlapping region. For every threshold calculated, the proportion of overlapping regions was larger in MS NAWM tissue (blue) than in non-neurological control tissue (grey). (F) The mean paranodal length per block correlated with the proportion of overlapping regions at an intensity threshold of 50 (r = 0.49, **p < 0.01, Spearman’s rank correlation test). (G) The proportion of paranodes longer than 4 μm per block correlated with the proportion of overlapping regions at a threshold of 50 (r = 0.57, ***p < 0.001, Spearman’s rank correlation test).

### Paranodal disruption is associated with juxtaparanodal K_v_1.2 channel dislocation

One of the roles of the tightly adherent axo-glial junctions is to promote the clustering and segregation of voltage gated channels and prevent electrical current shunting underneath the myelin sheath. To examine if paranodal structural instability could provoke a dislocation of juxtaparanodal voltage-gated K_v_1.2 channels to the paranode in MS NAWM tissue, RGB intensity profiles of the Caspr1 (red) and voltage gated K_v_1.2 channel (green) labelled paranodes and juxtaparanodes were measured (figure 2C,D). The co-localisation of Caspr1 and K_v_1.2 was calculated by subtracting both intensity signals one from each other. When this difference was smaller than a variable threshold, we took that point as an overlapping region (Supplementary Data). For a variable intensity threshold, MS NAWM tissue had a higher proportion of overlapping regions at every threshold than the non-neurological control tissue (figure 2E). For example, if the threshold was set to 50, the MS NAWM tissue had 19.7% more overlapping regions than non-neurological control tissue, whilst at a threshold of a 100, the MS NAWM tissue had 41% more overlapping regions than control. Thus, paranodal disruption was accompanied by a dislocation of the juxtaparanodal K_v_1.2 channels towards the node, making these voltage-gated channels more exposed to the extracellular space. Additionally, significant correlations were present between the proportion of overlapping regions in NAWM tissue when the threshold was set to 50 and the mean paranodal length (figure 2F) and proportion of paranodes longer than 4μm (figure 2G). The same quantification procedure was followed for assessing dislocation of the nodal Na_v_ voltage-gated channels into the PNJ, but no significant dislocation was identified (Supplementary Figure2).

### Prolonged exposure of the rat cortex to pro-inflammatory cytokines can generate paranodal disruption

To further study the relationship between chronic inflammation in the NAWM and PNJ pathology, we used a novel rat model of chronic meningeal inflammation (James et al, 2020). In this model, inflammation was initiated by chronic production of the pro-inflammatory cytokines LTα and IFNγ within the meninges and cerebrospinal fluid (CSF). Meningeal inflammation induced widespread microglial activation over 3 months in the absence of WM demyelination, which did not occur in these animals (Supplementary Figure3). This allowed the study of the effects of diffuse neuroinflammation on the axons of the NAWM, similar to those seen in progressive MS, avoiding Wallerian degeneration and dying back axonal injury patterns resulting from demyelination. Control groups were rats injected with a green fluorescent protein (eGFP) gene vector or naive rats (Supplementary Figure3). NAWM regions were selected from the MOG immunofluorescence images of the corpus callosum, cingulate and external capsule (figure 3A).

**Figure 3:**
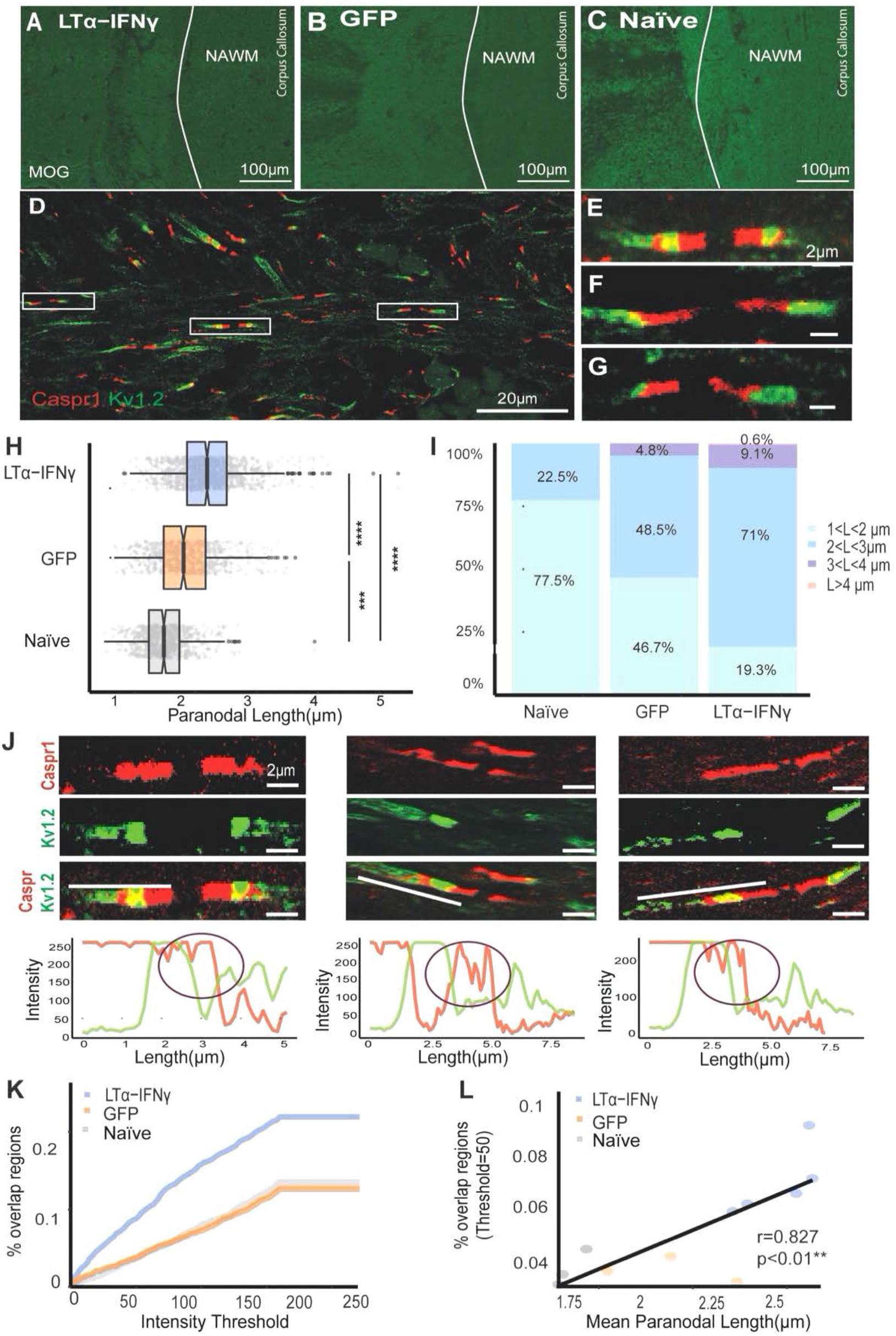
LTα/IFNγ rat tissue contained a higher proportion of elongated Caspr1-paranodes compared to the GFP and naive. (A,B,C) Immunofluorescent images of MOG stained corpus callosum, cingulate and external capsule of the LTα/IFNγ, GFP and naive rats. (D) Confocal image of a Caspr1-K_v_1.2 stained paranodes and juxtaparanodes from a LTα/IFNγ rat. (E,F,G) Confocal images of single Caspr1-K_v_1.2 stained paranodes and juxtaparanodes from a LTα/IFNγ rat. (H) Caspr1-measured paranodal length distributions of LTα/IFNγ (blue), GFP (orange) and naive (grey) rat groups (**** p< 0.0001, Mann-Whitney test). (I) Paranodal length data was divided into different length ranges and represented in a bar plot. LTα/IFNγ injected brains contained a larger proportion of paranodes longer than 3 and 4 μm. (J) Confocal images in which Caspr1-stained paranodes co-localised with K_v_1.2-stained juxtaparanodes. In order to quantify the displacement of the channels the RGB intensity profiles of each paranode and juxtaparanode were acquired. The purple circles denote regions where both signals were co-localising. (K) Graph showing that the proportion of overlapping regions between Caspr1 and K_v_1.2 RGB signals was larger at every intensity threshold in the LTα/IFNγ group compared to the GFP and naive groups. (L) Mean paranodal length correlated significantly with the proportion of overlapping regions when the intensity threshold was set to 50 (r=0.827, **p< 0.01, Spearman’s rank correlation test).

Immunofluorescence analysis of Caspr1 and voltage-gated K_v_1.2 channel localisation was carried out as for the human tissue (figure 3 D-G). In total, 1000 Caspr1-stained paranodes (200 paranodes per rat) from the LTα/IFNγ group, 600 from the GFP group, and 600 from the naïve group were analysed. The mean Caspr1 stained paranodal length in the LTα/IFNγ vector group (2.41 ± 0.01μm) was 35.6% longer than values for naive rats (1.77 ± 0.01μm) and 15.9% longer than the GFP-vector injected rats (2.08 ± 0.02μm). Furthermore, 82% of the paranodes in the LTα/IFNγ group were longer than the 75% percentile of the naïve group (1.98μm) (figure 3H). Paranodal length distributions of each group showed that 9.7% of the paranodes in the LTα/IFNγ group were longer than 3μm compared to the 4.8% in the GFP group and 0 % in the naïve group (figure 3I (purple and pink)). Compared to the naïve group, in which 77.5% of the paranodes were shorter than 1μm, in the GFP group 46.7% and in the LTα/IFNγ group 19.3% were shorter that 1μm in length (figure 3I). Thus, paranodal axo-glial junctions in the LTα/IFNγ vector injected animals, in which there was widespread microglial activation, were highly disrupted compared to the GFP and naïve groups.

Analysis of the RGB intensity profiles of Caspr1 and K_v_1.2 channels, following the same method as in the post-mortem human tissue (Supplementary Data), showed that the brains of LTα/IFNγ vector injected animals contained a larger proportion of axons with overlapping regions (figure 3J-K) at every threshold analysed. Therefore, the LTα/IFNγ animals had more damaged axons with dislocated voltagegated K_v_1.2 channels, indicating more disrupted PNJs than the GFP and naïve groups (figure 3J,K). When the intensity threshold was set to 50, the proportion of overlapping regions were significantly different between LTα/IFNγ and GFP or naïve groups. Moreover, the proportion of overlap correlated with the average paranodal length (figure 3L, r=0.827) and with the proportion of paranodes >3μm per rat (r=0.765). These results suggest that paranodal lengthening and K_v_1.2 dislocation were often observed together. To investigate whether surrounding inflammation could have a role in this pathology, the number of microglia and astrocytes in the NAWN regions were assessed on serial sections from the same animals (figure 4A-F). The number of microglia in the NAWM correlated with the average paranodal length (figure 4G, r=0.62) and with the proportion of overlapping regions between Caspr1 and K_v_1.2 (figure 4H, r=0.778). Furthermore, the number of astrocytes also correlated with the average paranodal length (figure 4I, r=0.69) and with the proportion of overlapping regions between Caspr1 and K_v_1.2 (figure 4J, r=0.75). This data supports our hypothesis that points to microglia and astrocytes as major mediators of the paranodal axo-glial junction disruption in myelinated axons.

**Figure 4:**
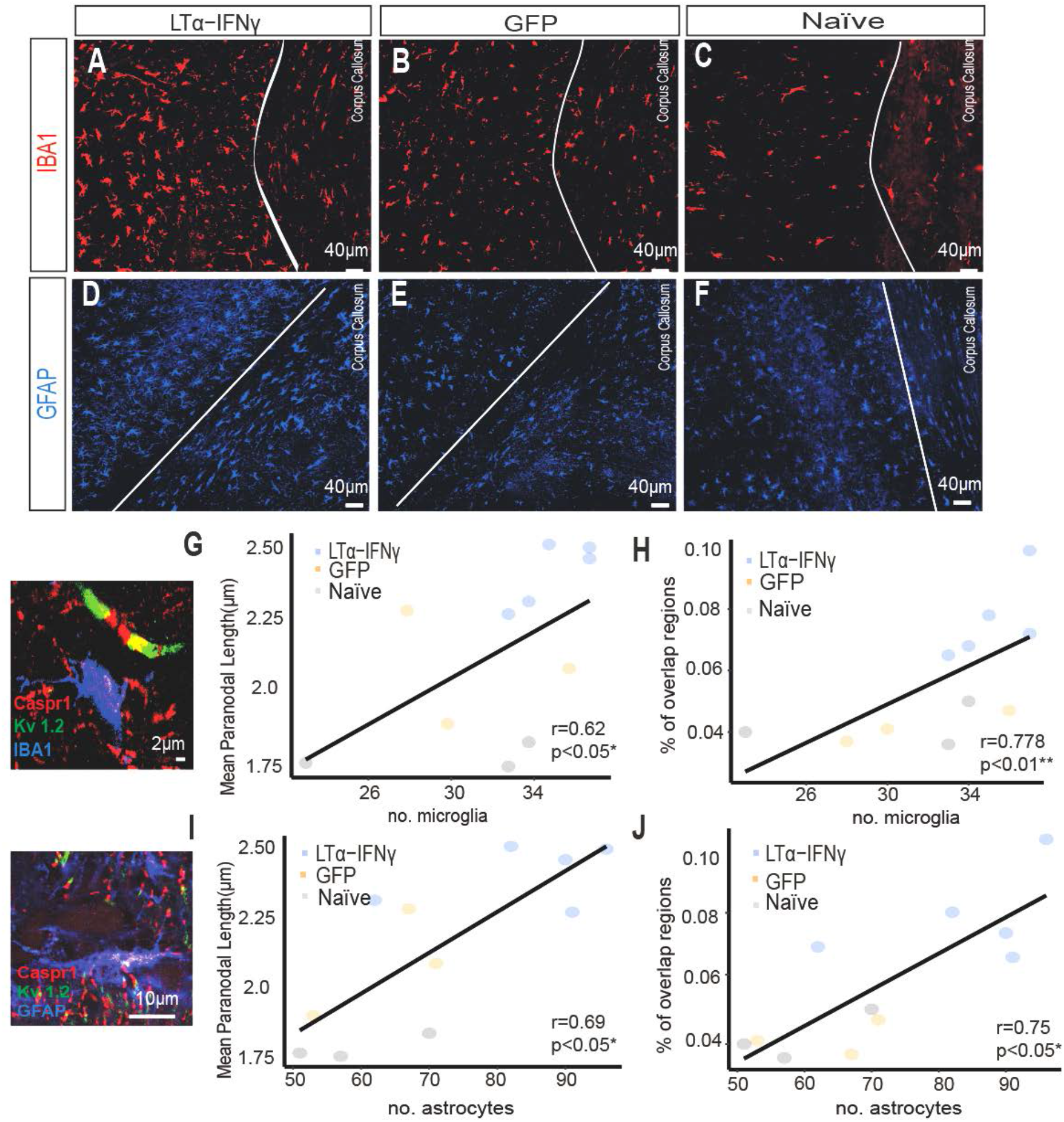
Paranodal lengthening and K_v_1.2 dislocation occurs simultaneously, and they are linked to glial activation. (A,B,C) Immunofluorescent images of IBA1+ microglia from the corpus callosum of LTα-IFNγ, GFP and naive rats. (D,E,F) Immunofluorescent images of GFAP+ astrocytes from the corpus callosum of the LTα/IFNγ, GFP and naive rats. (G) The number of microglia correlated with the mean paranodal length per rat (r=0.62, *p< 0.05, Spearman’s rank correlation test). (H) The number of microglia correlated with the proportion of overlapping regions between Caspr1 and K_v_1.2 when the intensity threshold was set to 50 (r=0.778, **p< 0.01, Spearman’s rank correlation test). (I) The number of astrocytes correlated with the mean paranodal length per rat (r=0.69, *p< 0.05, Spearman’s rank correlation test). (J) The number of astrocytes correlated with the proportion of overlapping regions between Caspr1 and K_v_1.2 when the intensity threshold was set to 50 (r=0.75, *p< 0.05, Spearman’s rank correlation test).

### TNF/IFNγ activated microglia release high levels of glutamate

Previous studies have suggested that elevated glutamate levels are able to induce nodal changes (Fu et al., 2009, 2011; Huff et al., 2011). Therefore, in order to assess if increased levels of pro-inflammatory cytokines in the CNS can stimulate glutamate release from microglia, primary rat microglial cultures were treated with different concentrations of TNF, IFNγ or TNF + IFNγ. Microglia were activated by TNF and/or IFNγ as indicated by transformation of their morphology from a ramified shape to a more activated amoeboid morphology (figure 5 B-D). A single treatment with 100 ng/ml of TNF, IFNγ or TNF+IFNγ, induced a significant increase in glutamate release after 24hrs (TNF: 88.68±26.67 μM, IFNγ: 81.62±10.4μM, TNF+IFNγ: 83.69±2.88 μM) and 48hrs (TNF: 79.39±7.64μM, IFNγ: 52.95±4.14 μM, TNF+IFNγ: 76.02±8.42 μM) compared to controls (24hrs - 33.49±2.11μM; 48hrs - 27.73 ±7.33μM)(figure 5 E), which did not increase further if the dose was increased to 200 ng/ml (Supplementary Figure 4). The difference in glutamate levels between 24hrs and 48hrs across treatments was not significant (figure 5E,F). However, administration of two doses of 100 ng/ml of TNF+IFNγ 24 hours apart (figure 5F) resulted in the highest concentration of glutamate in the medium at 48hrs (96.046±9.92 μM) when compared to the single treatments. We have used TNF and LTα interchangeably here as they both act via the TNFR1 receptor in this context. The same experiments on the primary microglial cultures were also carried out with the pro-inflammatory cytokines LTα and IFNγ, which also resulted in a significant glutamate release compared to controls after two doses administered 24 hrs apart (Supplementary Figure 4).

**Figure 5:**
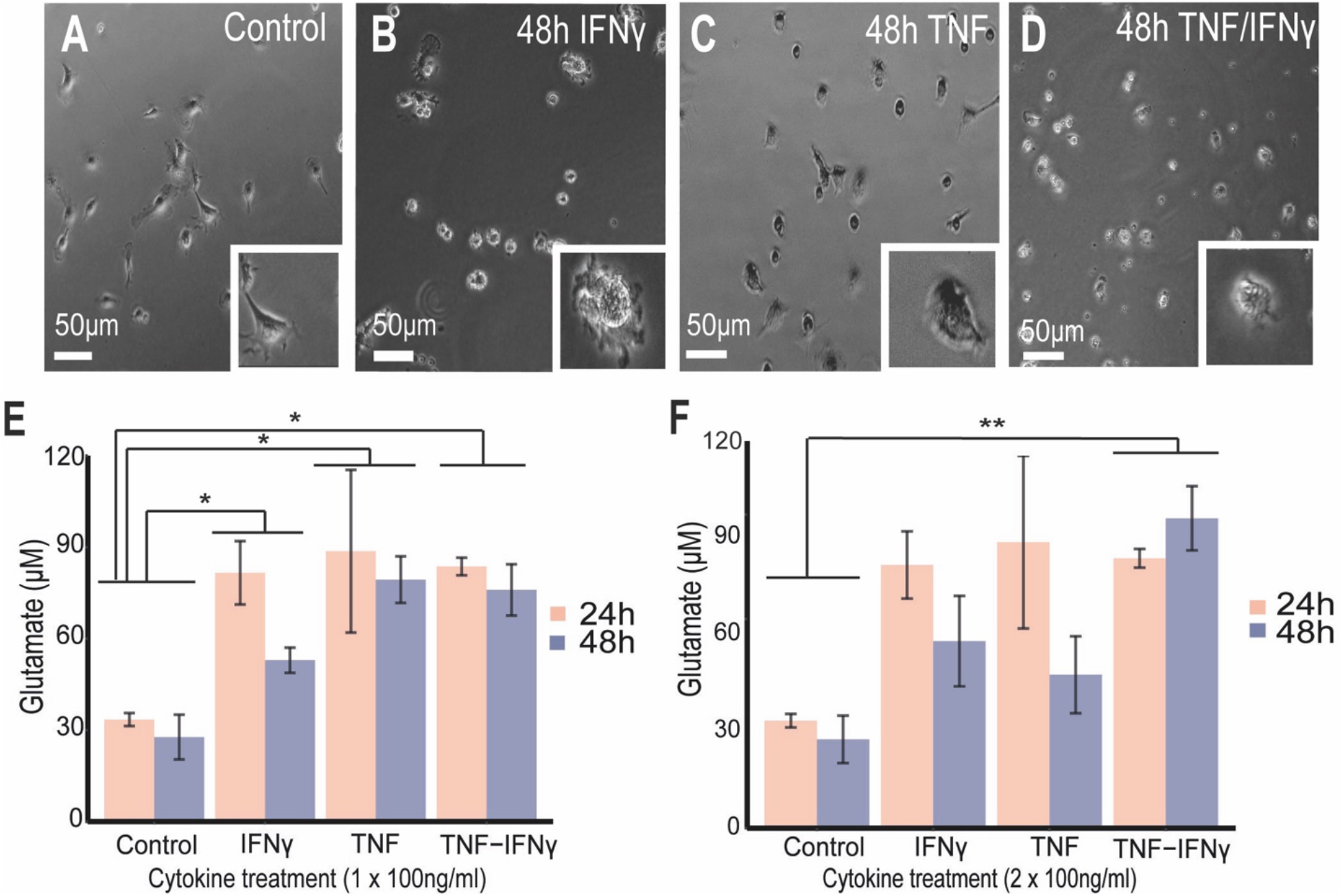
TNF/IFNγ activated microglia release high amounts of glutamate. (A) Live image of non-treated, (B) treated with IFNγ after 48hrs, (C) treated with TNF after 48hrs and (D) treated with TNF/IFNγ after 48hrs of primary microglia cultures. (E,F) Mean ± SEM for glutamate levels from replicates showing the statistical difference between controls and the cytokine treatments: (E) 100 ng/ml (n=3 Control, n=3 TNF, n=3 IFNγ, n=3 TNF/IFNγ), (F) two acute treatments of 100 ng/ml (n=3 Control, n=3 TNF, n=3 IFNγ, n=4 TNF/IFNγ). Non-parametric Friedman test was performed across cytokine groups and timings and post hoc paired-wised wilcoxon tests to compare groups (* p<0.05, ** p<0.01).

### TNF/IFNγ and direct glutamate administration induce paranodal elongation in ex-vivo cerebellar slices

In order to determine if TNF + IFNγ and glutamate could cause MS-like paranodal pathology, organotypic cerebellar slices derived from P8/9 rats and cultured for 8-10 days were treated with three doses of 50 ng/ml TNF/IFNγ, two doses of 100 ng/ml of TNF/IFNγ, two doses of microglial conditioned medium, two doses of glutamate at 75μM or two doses of glutamate at 100μM (in all the treatments doses were administered every 24hrs and glutamate levels were measured 24hrs after the last dose, Supplementary Figure 5). For each of the cerebellar tissue slices, 200 focused Caspr1-stained paranodes were analysed (figure 6A). In cerebellar slices treated with treated with TNF/IFNγ, Caspr1-stained paranodal length was 19.15% longer on average in the 50 ng/ml group (2.96 ± 0.03μm) and 16.6% on average in the 100ng/ml group (2.88 ± 0.04μm) than in the non-treated cerebellar cultures (2.47 ± 0.03μm, figure 6B). However, when separating the paranodal measurements by length ranges, they had larger proportions of highly disrupted long paranodes > 4μm than the untreated cerebellar slices (11.38% in the 100 ng/ml group and 11.34% in the 50 ng/ml group figure 6C, light and dark purple). Abnormally long Caspr1-stained paranodes were also observed (asterisks in figure 6A). Five cerebellar slices were treated with the microglial conditioned medium produced after two doses of 100ng/ml of TNF/IFNγ, the condition that resulted in the greatest glutamate release. These cerebellar cultures had an average paranodal length of 3.23 ± 0.04 μm, 32.84% longer on average than the non-treated ones (2.47 ± 0.027 μm; 200 paranodes were analysed per cerebellar slice, figure 6A-C). In order to examine if glutamate was one of the main cytotoxic factors in the microglial cytokine-conditioned medium, 4 cerebellar slices were treated with 75μM of glutamate for 48hrs (the average glutamate concentration in the 100ng/ml TNF/IFNγ group was 96.046 ± 9.92μM and the conditioned medium was diluted 4:1), which resulted in paranodes 26.11 % longer on average than the non-treated tissue slices (figure 6A-C).

**Figure 6:**
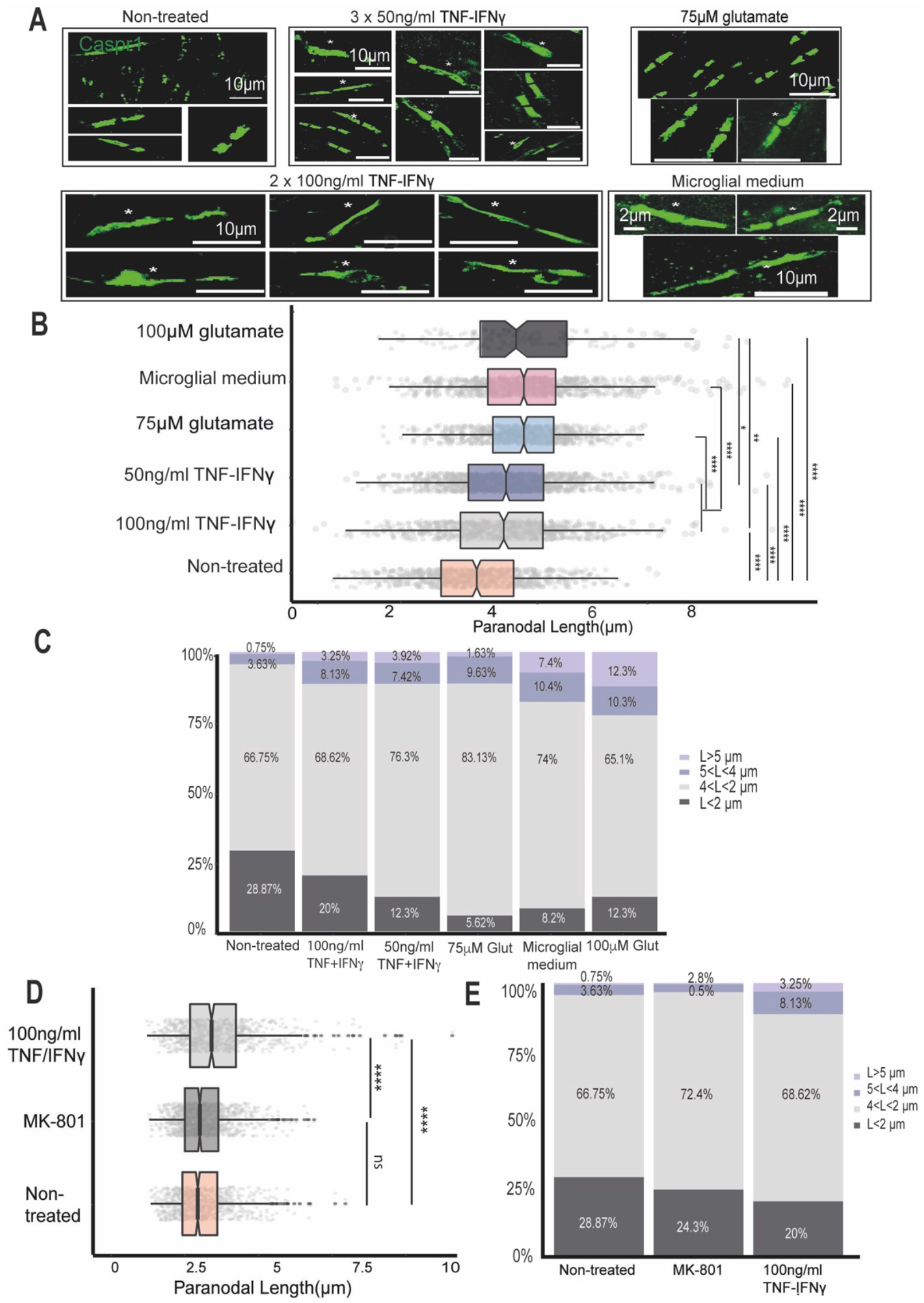
The proportion of elongated paranodes in cerebellar tissue slices treated with pro-inflammatory cytokines and glutamate. (A) Confocal images of Caspr1-stained paranodes from non-treated cerebellar cultures, slices treated with 3 doses of 50 ng/ml TNF/IFNγ, slices treated with 2 doses of 100 ng/ml TNF/IFNγ, slices treated with 2 doses of 75μM glutamate and slices treated with 2 doses of microglial-conditioned medium (medium from primary microglia treated with two doses of 100ng/ml TNF/IFNγ); asterisks point to long and disrupted paranodes. (B) Box-plots showing the different paranodal length distributions between the treated and non-treated cultures (non-parametric Kruskal Wallis test and post-hoc Wilcoxon rank-sum test, ****p< 0.0001). (C) Bar plots of the same paranodal length distributions showing the proportion of paranodes in each data set of different lengths. (D) Box-plot of the paranodal length distributions of the non-treated (orange), MK-801 (dark grey) and two doses of 100 ng/ml TNF+IFNγ cultures (**** p< 0.0001, Mann-Whitney test). (E) Bar plot showing the same paranodal length distributions divided into different length ranges.

In order to confirm if the effect of TNF/IFNγ was due to glutamate release and action, 5 slices were treated with two doses of 100 ng/ml of TNF/IFNγ together with the non-competitive NMDA antagonist MK-801 (0.6mM) (figure 6D). The paranodal length distribution of the slices was not significantly different between the MK-801 treated and non-treated slices (figure 6E). The proportion of paranodes longer than 4μm was greatly reduced in the MK-801 group compared to the cultures treated with the cytokines alone (from 11.38% of the paranodes to 3.3%, figure 6E).

### PNJ disruption can affect velocity and conduction in small-diameter axons

To systematically examine the consequences of paranodal disruption on AP propagation through the use of computational modelling, a double cable core model was built and solved numerically in NEURON (figure 7A). A double cable core-conductor circuitry was chosen to represent the biophysical parameters of the axonal membrane, the peri-axonal space and the myelin sheath, separately (Richardson et al., 2000; McIntyre et al., 2002). We simulated the functionality of an axon membrane made up of 4 types of compartments: nodes, paranodes, juxtaparanodes and internodes. The nodes clustered a high density of fast Na_v_ channels (Na_v_1.6), persistent Na_v_ channels and slow K_v_ channels (figure 7A, purple). Immediately flanking the nodes, the paranodes were built as compartments with no active conductances, and the sites where the myelin end loops connect with the axolemma (figure 7A, dark grey). Next to the paranodes, the juxtaparanodes contained fast K_v_ channels (figure 7A, medium grey). Finally, the internodes were sections surrounded by myelin with a low density of ion channels (figure 7 A, light grey). Seven fibre diameters were simulated: d _fibre_ = 0.5, 0.8, 1.1, 1.3, 1.6, 1.8, 3.5 [μm]. These small-caliber diameters were chosen taking into consideration previous human brain and macaque EM studies, which indicated that the average axon core diameter within the CNS is 1μm (Liewald et al., 2014). All the structural and biophysical parameters used in the simulations were taken from previous EM studies and are summarised in figure 7B,C, whereas the conductances and gating dynamics of the channels were based on previous electrophysiological studies, detailed in the Methods and Materials. The conduction velocity predicted by our simulations showed a linear relationship with the fibre diameter (V[m/s] = 4.52 * d_fibre_ [μm]) (figure 7 D, blue line), which reproduced previous experimental results (figure 8D, dotted grey line, velocity data derived by Boyd and Kalu (1979) from small diameter axons in the cat hind limb nerves V = 4.6 * d_fibre_).

**Figure 7:**
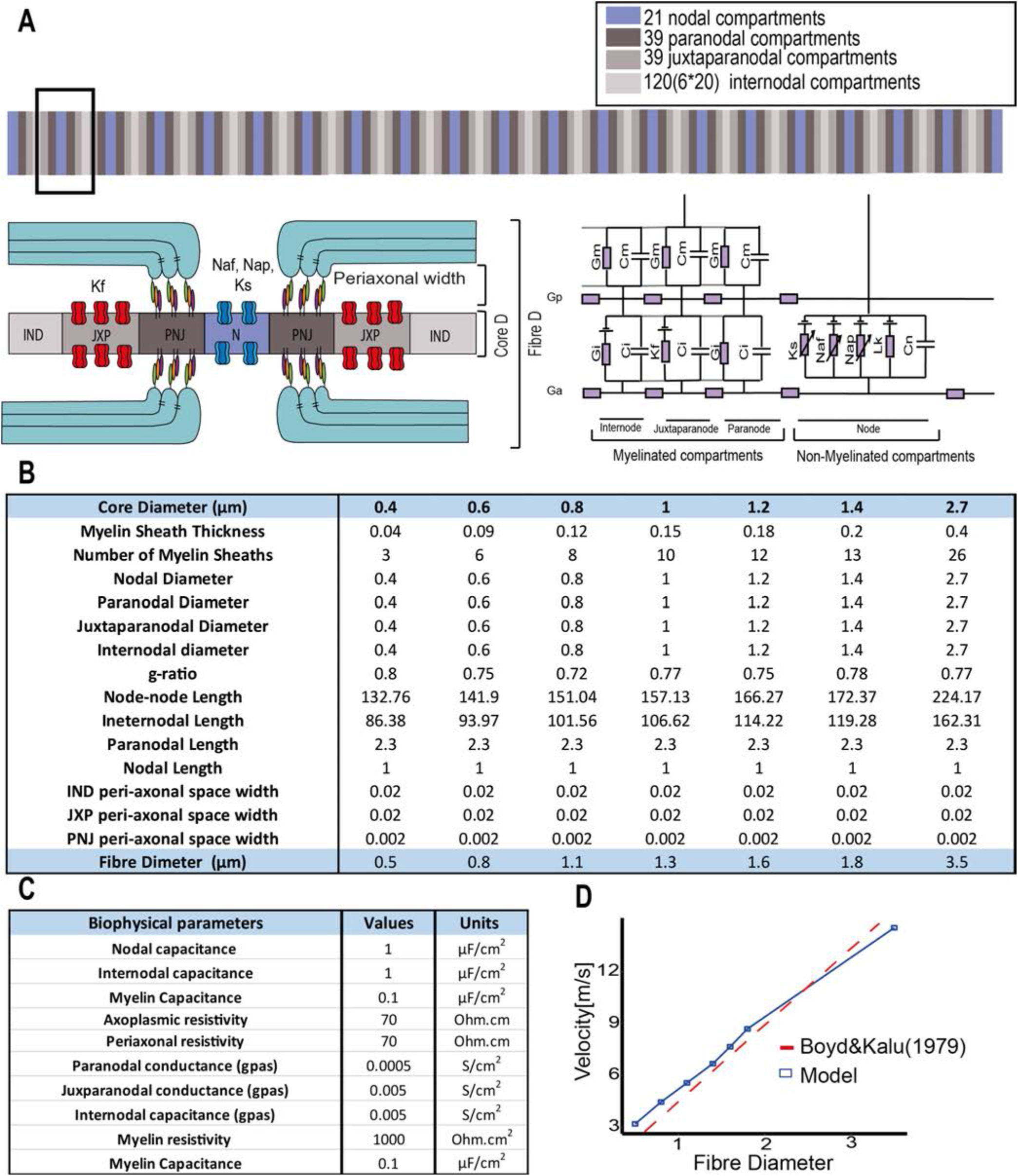
Structural and biophysical parameters used to simulate a 21 node myelinated axon. (A) A double cable circuit of the model to represent the axolemma (Ga), the peri-axonal space (Gp) and the myelin sheath (Gm) was generated with the simulator NEURON. Specifically, 21 nodal, 39 paranodal, 39 juxtaparanodal and 20 internodal compartments were created. (B) Anatomical parameters used in the model. Seven diameters were chosen from CNS measurements from macaque EM studies (Liewald et al., 2014), the number of myelin lamella (nl) was calculated from the myelin periodicity value of 0.0156μm (Agrawal et al., 2009), the node-to-node length was taken from the linear relationship measured from rat nerve fibres (Ibrahim et al., 1995), the juxtaparanodal length was extrapolated from the diameter dependent scaling relationship from the ventral root of cats (Berthold and Rydmark, 1983) and the paranodal length was determined from the average value of Caspr1 staining measured from our non-neurological control cases. (C) Biophysical parameters. Axon capacitance was based on data from rat ventral roots (Bostock and Sears, 1978), myelin capacitance and leak conductance per lamella were based on the frog sciatic nerves (Tasaki, 1955). The resistivity was set to 1000 Ohm * cm^2^ for each myelin lamella (Tasaki, 1955; Barrett and Barrett, 1982; Halter and Clark, 1991; McIntyre et al., 2002; Gow and Devaux, 2008; Boucher et al., 2012). (D) Plot showing the conduction velocity of the model across the seven fibre diameters simulated (blue) and the velocity data measured in cat hind limb nerves (Boyd and Kalu, 1979).

In order to simulate paranodal disruption, the resistance of the paranodal and/or juxtaparanodal compartments was decreased, and the juxtaparanodal K_v_ channels were dislocated. The resistance was decreased by increasing the peri-axonal space of both compartments progressively. The observed paranodal lengthening was represented in this model by an increment in peri-axonal space on the assumption that if some myelin-end loops at the PNJ detached from the axolemma, these spaces will be progressively larger (Bhat et al., 2001; Zonta et al., 2008). Furthermore, juxtaparanodal K_v_ channels were dislocated to the paranode, and their conductance was increased proportionally to the increment in peri-axonal space of the paranode. In summary, the following structural arrangements were simulated: (a) Proportional increment of the paranodal peri-axonal space (figure 8A); (b) proportional increment of the paranodal and juxtaparanodal peri-axonal space (figure 8B); (c) proportional increment of the juxtaparanodal fast K_v_ channels conductance and the paranodal peri-axonal space (figure 8C); and the proportional increment of the juxtaparanodal fast K_v_ channels conductance, and the paranodal and juxtaparanodal peri-axonal spaces (figure 8D).

**Figure 8:**
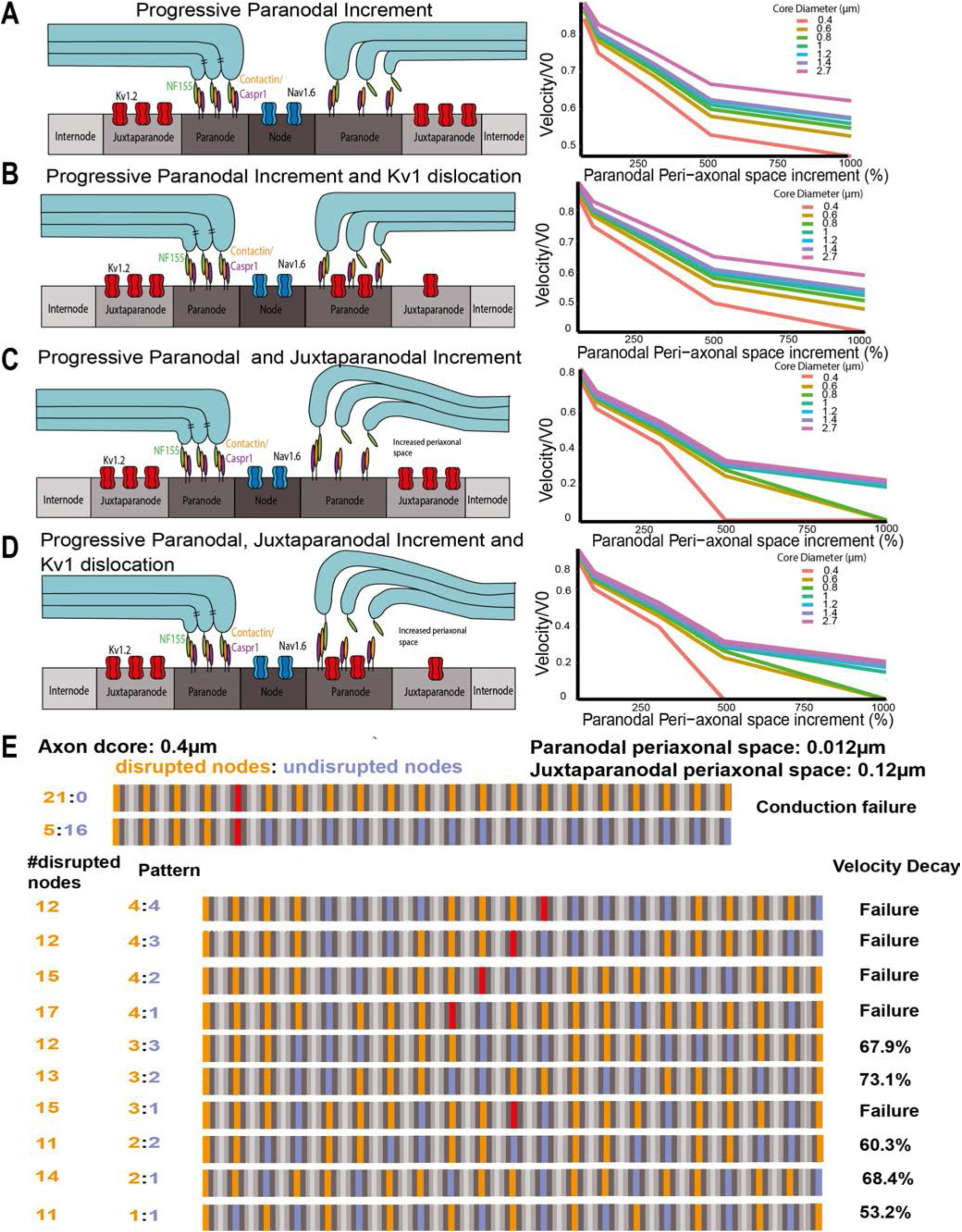
The effects of paranodal disruption on AP conduction is inversely proportional to the axon diameter. (A) Diagram representing the increment of the paranodal peri-axonal space and a normalised velocity plot at every core diameter as the paranodal peri-axonal space increases. (B) Diagram representing the increment of the paranodal and juxtaparanodal peri-axonal space and a normalised velocity plot at every core diameter as the paranodal peri-axonal space increases. (C) Diagram representing the increment of the paranodal and juxtaparanodal peri-axonal and K_v_1 dislocation, and a normalised velocity plot at every core diameter as the paranodal peri-axonal space increases. (D) Diagram representing the increment of the paranodal and juxtaparanodal peri-axonal and K_v_1 dislocation, and a normalised velocity plot at every core diameter as the paranodal peri-axonal space increases. (E) In an axon model with a core diameter of 0.4 μm, conduction failure occurred when five consecutive nodes were disrupted (orange) and the paranodal and juxtaparanodal peri-axonal space widths were increased up to 0.012 and 0.12 μm, respectively. Further, the velocity can decay and conduction can fail under different patterns of disruption (orange means disrupted node, purple, healthy node and, red denotes conduction failure.

The conduction velocity decreased as the paranodal peri-axonal space increased in all the axon diameters (figure 8A). Fitting the data to a logarithmic curve, the velocity of the axons decreased faster in the smaller-diameter axons than the larger-diameter ones as the paranodal peri-axonal space was increased (figure 8A). When the juxtaparanodal fast K_v_ channels were dislocated to the paranodal compartment (figure 8B), the velocity also decreased faster in the small-diameter axons, although the dislocation of the channels did not significantly alter this. When a progressive increment of the paranodal and juxtaparanodal peri-axonal spaces was introduced (figure 8 C), the axon with a core diameter of 0.4 μm failed to conduct when the paranodal and juxtaparanodal peri-axonal spaces were increased by 500% (the paranodal peri-axonal space was 0.012μm and the juxtaparanodal 0.12μm). Meanwhile, the axons with dcore of 0.6 μm and 0.8μm also failed to conduct when the paranodal and juxtaparanodal peri-axonal spaces were increased by 1000% (the paranodal peri-axonal space was 0.022 μm and the juxtaparanodal 0.22μm). Additionally, larger-diameter axons had a significant velocity decay in the same conditions. For example, the velocity of the axon with a core diameter of 2.7 μm decreased 78.47 % (y = −0.209 * ln(x) + 1.6457, r^2^= 0.99). In the last structural arrangement, the paranode and juxtaparanode peri-axonal spaces were increased and the K_v_1 were dislocated (figure 8 D). In this condition, conduction failure occurred in the axons with a core diameter of 0.4, 0.6, and 0.8 μm and a decrease in axons with a core diameter of 2.7 μm. This data suggests that AP conduction in the smaller-diameter axons might be more susceptible to paranodal and juxtaparanodal disruption than larger-diameter axons.

In the previous four conditions, all 21 nodes of the axon were disrupted by increasing the peri-axonal space widths and displacing the juxtaparanodal voltage-gated K_v_ channels. However, conduction failure only occurred in the small-diameter axons (d_core_ of 0.4, 0.6 and 0.8 μm). Therefore, we also examined the number of consecutive nodes needed for conduction failure at these diameters, and the velocity decay after simulating conduction in an axon where patches of healthy nodes (figure 8 E, purple) were interspersed with patches of disrupted nodes (figure 8E, orange), which is more likely to reflect the pathological situation. In the axon with d_core_= 0.4 μm, five consecutive nodes were needed for conduction failure when the width of the paranodal and juxtaparanodal peri-axonal spaces were increased to 0.012 and 0.12 μm respectively (figure 8E). We then explored if damaging less than five nodes consecutively in different patterns would also cause conduction failure or a decrease in conduction velocity (figure 8E). By simulating different patterns, we could observe that the total number of disrupted nodes along an axon did not determine if the conduction was going to fail, but it depended on the number of consecutive nodes disrupted. For example, when three disrupted nodes (figure 8E, orange) were followed by two healthy nodes (figure 8E, purple) the conduction did not fail, instead it caused a velocity decay of 73.1 %. Meanwhile, in other conditions with less numbers of disrupted nodes, the conduction failed when four consecutive nodes were disrupted (figure 8E). Simulations using different patterns for axons with d_core_= 0.6 and 0.8μm can be found in Supplementary Figures 6 and 7.

## Discussion

The nodes of Ranvier represent the only points of direct contact between myelin and the axon, specifically at the paranodal axo-glial junctions (Simons et al., 2014; Stassart et al., 2018), and it is suggested that these sites could be one of the targets of the immune mediated attacks in MS. Paranodal and juxtaparanodal pathology has been described in only a few studies in human MS tissue (Wolswijk and Balesar, 2003; Coman et al, 2006; Howell et al., 2006, 2010) and its effects and causes remained unknown. Our detailed analysis of the longitudinal elongation of the Caspr1-stained paranodal axo-glial junctions and dislocation of the K_v_1.2 channels indicate a clear disruption in a proportion of paranodes in MS NAWM compared to non-neurological control tissue. These findings were reproduced in both an in-vivo and an ex-vivo model by the presence of persistently elevated levels of TNF/LTα and IFNγ, which indicate that proinflammatory cytokines could be the trigger for this pathology. Further investigations using primary microglial cultures and organotypic cerebellar slice cultures suggested that glutamate release by microglia in response to stimulation with pro-inflammatory cytokines could mediate these pathological changes at the paranodes. Accumulation of abnormal nodes of Ranvier could be responsible for some of the generalised MS symptoms that cannot be attributed to focal lesions.

The elongation of paranodal axo-glial junctions elongation, indicated by the paranodal axonal protein Caspr1 or by its glial counterpart, Nf155, suggests that the glial and axonal proteins may have detached from each other, leading to diffusion along the axolemma. Therefore, it could represent a partial or complete detachment of the myelin end loops, and between the loops themselves, that could result in a complete detachment of the myelin tongues. Although we have not directly demonstrated this using ultrastructural analysis, conditional murine knockouts of the paranodal proteins Caspr1, Nf155, βII spectrin and 4.1.B (Bhat et al., 2001; Zonta et al., 2008; Zhang et al., 2013; Cifuentes-Diaz et al., 2011; Buttermore et al., 2011) showed a lack of tight septate junctions and an increased peri-axonal space at the ultrastructural level, which was associated with an elongation of the paranodal protein profiles and dislocation of the juxtaparanodal voltage-gated channels K_v_1 towards the PNJ, similar to that seen in MS tissue. This structural change would mean an increment in the peri-axonal space width in the paranode and juxtaparanode and, therefore, a progressive change in the membrane capacitance of these compartments. The disruption in the adequate anchorage of K_v_1.2 voltage-gated channels within the juxtaparanode has been shown in previous studies in which the expression of TAG1 and Caspr2, which anchor K_v_1 channels to the axolemma, appeared reduced in NAWM MS tissue (Kastriti et al., 2015). Thus, the partial to complete disruption of the paranode, as well as the possible dissociation of the tripartite juxtaparanodal complex of TAG-1/Caspr2/Kv1, could explain voltage-gated K_v_1 channel diffusion along the membrane. Overall, these histopathological results are especially relevant in progressive MS since paranodal pathology can affect the physiological integrity of axons and can lead to incremental deficits if the inflammatory stimuli persist.

The histopathological analysis of the human tissue showed that a significant proportion of Caspr1-immunopositive elongated paranodes within the MS NAWM tissue were associated with the activated microglia surrounding them. It has been suggested that diffuse axonal injury in the NAWM is closely associated with activated microglia (Kutzelnigg et al., 2005; Moll et al., 2011; Singh et al., 2013) and we can now add paranodal pathology as a characteristic of the NAWM changes. The relationship between neuroinflammation and paranodal disruption in myelinated axons was further consolidated by our rat model of meningeal inflammation induced by the chronic expression of LTα and IFNγ. Previous studies have demonstrated the presence of paranodal elongation in NAWM regions in a mouse EAE model (Howell et al., 2010). However, in this model the changes in the NAWM were accompanied by the presence of focal demyelinating lesions and occurred over a relatively short time period. On the contrary, the model used for the current experiments was characterised by widespread microglial activation caused by the chronic presence of pro-inflammatory cytokines in the CSF and the absence of demyelinating lesions within the brain. The longitudinal elongation of the Caspr1 epressing paranodal axo-glial junctions and dislocation of the voltage-gated K_v_1.2 channels demonstrated that paranodal disruption can occur in the absence of demyelination. Therefore, if no anterograde and/or retrograde axonal degeneration is contributing to the observed paranodal disruption, focal inflammatory mediators released by microglia and astrocytes surrounding the axons are likely to play an essential role. In fact, the extent of both the paranodal elongation and dislocation of K_v_1.2 channels correlated with the number of microglia and astrocytes surrounding the altered nodes.

One of the mechanisms of glial injury at these sites could be the disruption of the axo-glial transport along the myelinic channels of proteins that maintain the adequate structure and function of these junctions (Nave, 2010; Stassart et al., 2018). Alternatively, the changes to the structure could be caused by the activation of calcium-sensitive proteinases, such as Calpain1, if glutamate levels are not kept at homeostatic levels. Glutamate released along hemichannels by activated microglia could diffuse into the paranodal and juxtaparanodal peri-axonal spaces and generate paranodal pathology (Fu et al., 2009) by acting on NMDA receptors in the cytoplasmic channels of the myelin sheath, which include the PNJ and the adaxonal glial membrane (Nave, 2010; Velumian et al., 2011). Our data suggests that activated microglia release elevated amounts of glutamate in response to stimulation by pro-inflammatory cytokines. The effects of these pro-inflammatory cytokines are highly relevant in the context of MS as recent studies have shown that TNF, IFNγ and LTα levels are increased in the CSF of progressive MS patients and correlated with meningeal inflammation, cortical demyelination and activation of MHCII+ microglia (Magliozzi et al., 2010; 2018). In addition, increased CSF levels of TNF and IFNγ can induce endogenous expression of TNF and IFNγ in the underlying cortical parenchyma (James et al, 2020). TNF and LTα interacting with the TNFR1 receptor can induce induce glutamate release by microglia via a gap junction/hemichannel system in an autocrine manner (Kuno et al., 2005; Takeuchi et al., 2006;

Yawata et al., 2008) and inhibit glutamate transporters in astrocytes by inducing downregulation of EAAT2/GLT-1 mRNA (Chao and Hu, 1994; Wang et al., 2003; Sitcheran et al., 2005). The increment in length of Caspr1+ paranodes in the organotypic slices treated with TNF and IFNγ confirmed the significant role of these cytokines in the paranodal axo-glial pathology. The generation of a similar pathology by the direct administration of glutamate and its inhibition by the non-competitive NMDA antagonist MK-801, implicates glutamate signaling as a key molecular mediator driving paranodal pathology in the NAWM of the MS brain. This novel result also opens the possibility of therapeutic intervention by reducing microglial glutaminase activity, blocking microglial hemichannels (Cx32, Cx36, and Cx43), or by inhibiting TNF and IFNγ signaling.

Our biophysical model simulations of a CNS myelinated axon suggest that the functional consequences of changes at the paranode on AP conduction could be highly detrimental, even if only a small proportion of paranodes throughout a myelinated axon are disrupted. Recent computational models have simulated paranodal retraction by increasing the nodal width (Babbs and Shi, 2013), equating the nodal resistance to the paranodal resistance (Volman and Ng, 2014) or by reducing the number of myelin loops in the paranode (Kohan et al., 2017). However, these models only simulated these conditions in a single axon diameter (Babbs and Shi, 2013; Volman and Ng, 2014), or in a few of them (Kohan et al., 2018), and in axons that were usually ≥ 1μm. In our simulations, paranodal disruption was progressively generated by an increment in the peri-axonal space width, which led to velocity reduction and conduction failure in small diameter axons. This is in line with thin axon diameter biophysical theory on saltatory conduction with ion channel delocalization (Neishabouri & Faisal, 2014). Our data imply that, because axons with small diameter have a lower myelin sheath resistance and a higher axoplasmic resistance, they have a higher dependency on the structural integrity of the PNJ to shunt electrical currents and maintain fast and efficient conduction. Furthermore, these results also highlight the importance of modelling the paranodal and juxtaparanodal compartments, as well as the nodal and internodal compartments, as they are highly sensitive regions that are essential in balancing charge flows with the node of Ranvier. In MS NAWM tissue, the distribution and density of activated microglia and macrophages can vary significantly throughout the WM tracts of the brain and spinal cord. Taking this heterogeneity into consideration, we found that conduction failure occurred in axons with a fibre diameter smaller than 0.8μm with just 5 consecutive disrupted paranodes. Different patterns of disrupted and healthy paranodes made conduction fail at some point along the path in axons with a fibre diameter smaller than 1.1μm.

In conclusion, our data strongly suggests that diffuse pathology in the NAWM, which includes paranodal disruption, could be caused by the presence of local cytokine induced inflammation leading to excess glutamate release from microglia. Such paranodal pathology, that cannot be attributed to focal demyelinating lesions, would be expected to alter the efficiency and velocity of AP conduction, adding to the overall neurological dysfunction in MS. This could also be relevant to white matter changes seen in other neurodegenerative conditions in which chronic microglial activation is a feature.

## Materials and Methods

### Human post-mortem tissue

Tissue blocks for this study were provided by the UK Multiple Sclerosis Society Tissue Bank at Imperial College London. Brains were collected following fully informed consent via a prospective donor scheme following ethical approval by the National Multicentre Research Ethics Committee (MREC 02/2/39). Thirty-four tissue blocks (2×2×1 cm) were selected from twenty cases of neuropathologically confirmed SPMS (Table 1; 11 females) with a mean age of 54.3 yrs (range 38-76 yrs), mean post-mortem delay (PMD) of 22 hrs (range 12-48 hrs), and a mean disease duration of 28.4 yrs (range 12-42 yrs). Sixteen blocks were also selected from nine non-neurological control brains (Table 2; 3 females), with a mean age of 73.7 yrs (range 50-88 yrs) and a mean PMD of 23 hrs (range 8-48 hrs). All tissue blocks were from the pre-central gyrus (containing the primary motor cortex) and the cerebral peduncle located in the midbrain, due to the presence of white matter (WM) tracts that contain highly longitudinally aligned axons that would make quantitative analysis more precise. The blocks were fixed in 4% paraformaldehyde in PBS (4% PFA), cryoprotected in 30% sucrose, frozen in isopentane on dry ice, and stored at −75°C. Cryosections were cut at a thickness of 10μm and stored at −75°C.

**Table 1:** Demographic data for post-mortem MS cases. Age at death, post-mortem delay, gender and duration of disease of the MS cases used in this study. The case numbers represent the UK MS Society Tissue Bank case identifiers. The cause of death is that presented on the death certificates.

| Case No | Age (yrs) | PMD (hrs) | Gender | Duration (yrs) | Cause of death |
| --- | --- | --- | --- | --- | --- |
| MS404 | 55 | 17 | F | 34 | Pneumonia, septicaemia |
| MS406 | 62 | 23 | M | 42 | Pneumonia, MS |
| MS411 | 61 | 24 | M | 29 | Pneumonia, MS |
| MS422 | 58 | 25 | M | 25 | Pneumonia, MS |
| MS444 | 49 | 18 | M | 20 | Renal failure |
| MS461 | 43 | 13 | M | 21 | Bronchopneumonia |
| MS466 | 65 | 25 | F | 36 | Pneumonia, Lung carcinoma |
| MS478 | 63 | 24 | F | 39 | Bowel cancer, MS |
| MS500 | 50 | 32 | M | 29 | Urinary sepsis |
| MS510 | 38 | 19 | F | 22 | Pneumonia |
| MS517 | 48 | 12 | F | 25 | Chest sepsis, MS |
| MS523 | 63 | 20 | F | 32 | Bronchopneumonia, MS |
| MS530 | 42 | 15 | M | 21 | MS |
| MS535 | 65 | 12 | F | 40 | MS |
| MS542 | 76 | 12 | F | 35 | Pneumonia, MS |
| MS549 | 50 | 8 | M | 29 | MS |
| MS567 | 45 | 48 | F | 23 | MS |
| MS584 | 42 | 26 | F | 12 | MS |
| MS585 | 53 | 27 | F | 27 | Bronchopneumonia, MS |
| MS587 | 58 | 36 | F | 20 | Cardiac arrest, MS |
| **N=20** | **mean = 54** | **mean = 22** | **22F:12M** | **mean = 28** |  |

**Table 2:** Demographic data for post-mortem non-neurological control brains. Age at death, post-mortem delay and gender of the control cases used in this study. The case numbers represent the UK MS Society Tissue Bank (eg C48) and UK Parkinson’s Disease Brain Bank case identifiers (eg PDC29). The cause of death is that presented on the death certificates.

| Case No | Age (yrs) | PMD (hrs) | Gender | Cause of death |
| --- | --- | --- | --- | --- |
| C48 | 68 | 10 | M | Colon cancer |
| C54 | 66 | 16 | M | Not available |
| C72 | 77 | 26 | M | Pneumonia |
| C74 | 84 | 22 | F | Old age |
| C75 | 88 | 8 | M | COPD |
| C76 | 87 | 31 | M | Pneumonia |
| PDC29 | 82 | 48 | M | Liver and lung cancer |
| PDC39 | 50 | 28 | F | Renal cancer |
| PDC40 | 61 | 15 | F | Ovarian cancer |
| **N=9** | **mean = 74** | **mean = 23** | **5F:11M** |  |

### Primary and secondary antibodies

The primary antibodies used in this project were: mouse anti-MOG (clone Y10, Prof Reynolds, Imperial College London, UK); rabbit anti-myelin basic protein (MBP) (Polyclonal, Merck, Darmstadt, Germany); mouse anti-neurofilament-H protein (clone NE14; Merck, Darmstadt, Germany); mouse anti-dephosphorylated neurofilament protein (clone smi32; Biolegend, San Diego, CA, USA); rabbit anti-Caspr1 (clone EPR7828, Abcam, Cambridge, UK); mouse anti-Caspr1 (clone K65/35; Neuromab, Davis, CA, USA); mouse anti-pan-Na_v_ channels (clone K58/35, Neuromab, Davis, CA, USA); mouse anti-K_v_1.2 channels (clone K14/16, Neuromab, Davis, CA, USA); mouse anti-HLA-DR (clone TAL.1B5, Dako Agilent, Santa Clara, CA, USA); rabbit anti-IBA1 (Polyclonal IgG, Wako Pure Chemical Corporation, USA); rabbit anti-Glial Fibrillary Acidic Protein (GFAP) (Polyclonal, Dako Agilent, Santa Clara, CA, USA); mouse anti-Calbindin1 (clone CB-955, Merck, Darmstadt, Germany); mouse anti-NeuN (clone A60, Merck, Darmstadt, Germany) (Table 3). All the secondary antibodies used for immunofluorescence were purchased from ThermoFisher Scientific (USA): Alexa Fluor 546 Goat Anti-Mouse IgG (H+L), Alexa Fluor 546 Goat Anti-Rabbit IgG(H+L), Alexa Fluor 488 Goat Anti-Mouse IgG (H+L) Fluor 488 Goat Anti-Rabbit IgG (H+L), Alexa Fluor 488 Goat Anti-Mouse IgG1, Alexa Fluor 546 Goat Anti-Mouse IgG1, Alexa Fluor 647 Goat Anti-Mouse IgG1, Alexa Fluor 488 Goat Anti-Mouse IgG2b, Alexa Fluor 647 Goat Anti-Mouse IgG2b, Alexa Fluor 488 Streptavidin, Alexa Fluor 546 Streptavidin, Alexa Fluor 647 Streptavidin. For immunohistochemistry the following biotinylated antibodies were used from Vector Laboratories (UK): biotinylated anti-rabbit IgG (H+L) made in goat, biotinylated anti-mouse IgG (H+L) made in horse and biotinylated anti-mouse IgG2a made in goat (Life Technologies, ThermoFisher Scientific).

**Table 3:** Primary antibody details. Antigen, specificity, concentration, clone and source of the primary antibodies used in this project.

| Antibody | Specificity | Dilution | Clone | Source |
| --- | --- | --- | --- | --- |
| MOG | Myelin oligodendrocyte glycoprotein | 1:50 | Mouse (IgG2a) Y10 | Reynolds, Imperial |
| MBP | Myelin basic protein | 1:500 | Rabbit polyclonal | Merck |
| NFil | 200kD neurofilament protein | 1:500 | Mouse (IgG1) NE14 | Merck |
| SMI32 | De-phosphorylated neurofilament proteins | 1:500 | Mouse (IgG1) SMI32 | BioLegend |
| Caspr1 | Contactin-associated protein 1 | 1:300 | Rabbit EPR7828 | Abcam |
| Caspr1 | Contactin-associated protein 1 | 1:500 | Mouse (IgG1) K65/35 | Neuromab |
| Pan-Na<sub>v</sub> | Voltage gated sodium channels | 1:100 | Mouse (IgG2a) K58/35 | Neuromab |
| Kv 1.2 | Voltage gated potassium channel 1.2 | 1:100 | Mouse (IgG2a) K14/16 | Neuromab |
| HLA-DR | MHC class II human leucocyte antigen DR | 1:200 | Mouse TAL.1B5 | Dako Agilent |
| IBA1 | Ionized calcium binding adaptor molecule 1 | 1:500 | Rabbit polyclonal | Wako |
| GFAP | Glial fibrillary acidic protein | 1:500 | Rabbit polyclonal | Dako Agilent |
| Calbindin1 | Calbindin1 (Purkinje neurons) | 1:500 | Mouse (IgG1) CB-955 | Merck |
| NeuN | Neuron specific nuclear protein | 1:500 | Mouse (IgG1) A60 | Merck |

### Immunostaining on post-mortem human tissue

For immunofluorescence for all antigens, sections were fixed with 4% PFA for 30 min (Table 3), except for the Pan-Na channel antibody for which fixation was not longer than 10 min. After fixation, sections were post-fixed with 100% methanol at −20°C (Sigma) for 10min and washed in 0.1M PBS-0.3% Triton X-100 (Sigma-Aldrich) three times, five minutes each. After post-fixing, sections were blocked and permeabilized with 0.1 M PBS containing 10% normal horse/goat serum (Sigma-Aldrich) and PBS-0.3% Triton X-100 for 1 hr at room temperature. Finally, sections were incubated overnight at 4°C in a humid chamber with primary antibodies (Table 3) in 0.1 M PBS containing 10% normal horse/goat serum and PBS-0.3% Triton X-100 (Sigma-Aldrich). After primary antibody incubation, sections were thoroughly rinsed in 0.1M PBS at least three times, five minutes each. After rinsing, sections were incubated with the appropriate secondary antibody conjugated to biotin for 1hr for MOG, SMI32 and Pan-Nav antigens, and rinsed in 0.1M PBS. After rinsing, sections were incubated with Alexa Fluor Streptavidin or the appropriate species-specific secondary fluorochrome conjugated antibody, for 2 hrs at room temperature. Finally, the tissue was rinsed in 0.1M PBS and dH_2_O, nuclei counterstained with DAPI (diluted in 1:2000, Sigma-Aldrich) and mounted with Vectashield Antifade Mounting Media (Vector Laboratories). The coverslips were fixed to the slides with clear nail-polish.

Immunohistochemistry was performed for HLA-DR antigen with the ImmPRESS^TM^ HRP Anti-Mouse IgG (Peroxidase) Polymer Detection Kit, made in horse (Vector Laboratories). Tissue endogenous peroxidase activity was blocked with the Bloxall Blocking solution (SP-6000) for 10 minutes and then incubated with 2.5% horse serum for 20 min. After blocking and washing with 0.1M PBS, the primary antibody was incubated overnight at 4°C in a humid chamber. The ImmPRESS Polymer Reagent was incubated at room temperature for 30 minutes and the signal was developed with ImmPact DAB (SK-4105, Vector Laboratories). The tissue was then rinsed with tap water for 5min to stop the reaction. Slides were counterstained with haematoxylin (Sigma-Aldrich) for 5 min, washed with tap and distilled water, dehydrated with washes of 70%, 90% and 100% of ethanol (2 min each), cleared with xylene for 10min and mounted with DPX mounting medium (Sigma-Aldrich).

### Lentiviral vector production

Lentiviral (LV) vectors expressing the human LTα (LVLTα), human IFNγ (LVIFNγ) or enhanced green fluorescent protein (LVGFP) genes were produced exactly as described previously (James et al., 2020), using the human immunodeficiency virus type 1 (HIV-1) transfer vector (pRRL-sincppt-CMV-eGFP-WPRE genome plasmid) with a human cytomegalovirus promoter (CMV) promotor. Complementary DNA sequences (cDNA) for human LTα or IFNγ were codon optimised for rat, including a 5’ Kozak sequence, and synthesised by Gene Art (Life Sciences, Paisley, UK). The biological and physical titres of the purified and concentrated vectors were calculated as described previously (James et al., 2020). The lentiviral genome copy number was calculated using the Clontech Lenti-X qRT-PCR Titration kit (Takara).

### Stereotaxic surgery and tissue processing

All animal experiments were carried out under the regulations of the UK Home Office. Eleven 8-10 week old female Dark Agouti (DA) rats (140-160g) were obtained from Janvier (France) and kept in groups of 3-4 in a 12h light/dark cycle with food and water provided ad libitum. Stereotaxic injections of lentiviral vectors into the subarachnoid space were carried out at 0.9 mm caudal to bregma, in the midline, following previously published methods (Gardner et al, 2013; James et al., 2020). Rats were either naive with no treatment or were injected once with incomplete Freund’s Adjuvant (IFA). Rats were anaesthetised (2% isoflurane; Abbott Laboratories, Berkshire, UK and oxygen 2 l/min), their scalps were shaved and disinfected with Videne antiseptic solution (Ecolab). Subcutaneous injections of 0.9% saline (Sigma) and 0.01 mg/kg buprenorphine (Vetergesic; Alstoe Animal Health, North Yorkshire, UK) were performed to provide post-surgery rehydration. Rats were positioned on the stereotaxic frame (Stoelting, Dublin, Ireland) and an incision was made through the scalp to visualise the skull bregma. A 2mm diameter hole in the skull was drilled at the midline at 0.9mm caudal to bregma, at the level of the motor cortex. Injections were performed with a finely calibrated glass capillary attached to a 26-gauge needle of a 10 μl Hamilton syringe (Hamilton, Graubunden, Switzerland). The needle was inserted to a depth of 2.4 mm below the dural membrane. Four μl of lentiviral vector preparation (2 μl of each vector), diluted in TSSM with 0.5 mM monastral blue tracer, was introduced at a rate of 0.2 μl/min using an automated infuser (KD Scientific, USA). Viruses were injected at a total of 1×10^12^ genomic copies/μl (GC/μl) for LTα and GFP and 1×10^10^ GC/μl for IFNγ. The needle was left in place for 5 minutes to allow diffusion of the sample from the area of injection, then withdrawn and the incision was sutured (Mersilk; Covidien, Ireland). The sutures were removed after 7-10 days and the animals were monitored daily.

At the termination of the study, rats received an overdose of sodium pentobarbital (200 mg/ml Euthatal; Merial Animal Health, Essex, UK) by intraperitoneal injection. Rats were perfused with 50 ml PBS followed by 100 ml 4% PFA in PBS through the left ventricle at 90 days post vector injection. Brains were removed and post-fixed in 4% PFA (4 hours, room temp), prior to cryoprotection in 30% sucrose solution in PBS (48 hours or until equilibrated). Brains were embedded in optimal cutting temperature compound (OCT; Tissue-Tek; Sakura, The Netherlands), frozen in isopentane on dry ice and sectioned at 10μm in the coronal plane throughout the brain. The rat tissue sections were stained following the same protocol as the human tissue except the initial 4 % PFA fixation step.

### Primary rat microglial culture

P0-P2 Sprague Dawley rats (Charles River Laboratories, USA) were decapitated following the UK Home Office regulations. The brains were isolated, the cerebral cortices were dissected with sterile autoclaved tweezers and dissecting scissors and freed of meninges to avoid any fibroblast contamination. The cortices of three pups were transferred to a 50ml sterile polypropylene conical tube with dissection medium, which contained Hank’s Balanced Salt Solution (HBSS 1X, Gibco, ThermoFisher Scientific). The supernatant was removed and replaced by digestion mix containing Minimum Essential Media (MEM with 4 mM glutamine, Gibco, ThermoFisher Scientific), 2.5 mg/ml Papain (14 units/mg, Sigma-Aldrich, Merck), 40 g/ml DNAse (980 Units/mg, Sigma-Aldrich, Merck) and 240 g/ml l-Cysteine (Sigma-Aldrich, Merck). The 50ml Falcon tubes were placed in a water bath (Fisher Scientific) at 37°C for one hour; every 15 min the tissue was dissociated by gentle pipetting. After 1 hour the cortices were re-suspended in 30 ml warm dissection medium, filtered through a 70 μm nylon cell strainer (Corning Incorporated, USA) to remove any non-dissociated fragments and centrifuged at 500rpm for 5 min. The supernatant was discarded and the pellet was resuspended in 15ml of culture medium. The culture medium was Dulbecco’s Modified Eagle Medium Nutrient Mixture F-12 (DMEM F12, Gibco, ThermoFisher Scientific) supplemented with 5ml streptomycin-penicillin (10,000 U/mL, Gibco, ThermoFisher Scientific), 5ml L-glutamine (200 mM, Gibco, ThermoFisher Scientific) and 50 ml heat-inactivated fetal bovine serum (Sigma, Merck). The mixture of dissociated cells was plated in a T75 culture flask (Nunc EasyFlask 75cm^2^ cell culture flasks, Thermo Scientific), previously coated with poly-D-lysine hydrobromide (PDL, Sigma, Merck) for 2 hours and washed with sterile water 3 times. The mixed glial cultures were maintained for one week in a humidified incubator at 37°C and 5% CO2 for one week and half the medium was changed every 2 days to replace growth factors. After a week, astrocytes formed a connected confluent dense monolayer in the bottom of the plate, whereas the majority of the microglia were floating. Primary microglial cells were isolated from the astroglial cell bed by mechanical agitation by vigorously tapping the flasks. Subsequently, the cells were plated in a 24-well plate (Nucleon Delta Surface, Thermo Scientific) at a cell density of 5×10^4^ cells/well. After 24hrs, microglia were attached to the bottom of the plate and they were treated with the pro-inflammatory cytokines.

### Cerebellar organotypic tissue cultures

P8/P9 Sprague Dawley rats (Charles River Laboratories) were decapitated following the UK Home Office regulations. The hemispheres were separated and plated in a petri dish containing a sterile filter paper and cold dissection medium (Dulbecco’s Modified Eagle Medium (DMEM), supplemented with 5ml of streptomycin-penicillin (10,000 U/mL, Gibco, ThermoFisher Scientific). Hemispheres were mounted with Superglue together with a rectangular piece of sterile solid agar (4g of agar (Sigma) in 200ml of distilled water) and sliced using a Leica VT1200s vibratome in the parasagittal plane at 400μm thickness. The cerebellar slices were transferred into a 60mm petri dish containing cold medium until the whole hemisphere was sliced. Later, they were plated on sterile Biopore PTFE membranes of 0.4μm pore size (Millicell-CM culture inserts, Merck) in a sterile six well plate (Costar, Corning Incorporated) with 1 ml of nutrient medium underneath every culture insert. The nutrient medium contained 200ml Neurobasal-A-medium (Gibco, ThermoFisher Scientific) and 100ml Hank’s Balanced Salt Solution (HBSS 1X, Gibco, ThermoFisher Scientific) supplemented with 5ml streptomycin-penicillin (10,000 U/mL, Gibco, ThermoFisher Scientific), 5ml L-glutamine (200mM, Gibco, ThermoFisher Scientific), 4.4ml D-glucose (200 g/L, Gibco, ThermoFisher Scientific) and 2ml vitamin B27 Plus Supplement (50X, Gibco, ThermoFisher Scientific). One-two cerebellar slices were plated per insert and maintained in an incubator (HeraCell Vios 160i, ThermoFisher) at 37°C, 5% CO2 for 9-10 days, replacing half the medium every other day to replace the growth factors. In order to check the integrity of the slices, the macroscopic structure was checked with an inverted microscope (Olympus CKX53). They were also stained with the cell integrity marker propidium iodide (PI, Molecular Probes). PI was added at a concentration of 5 μg/ml, incubated for 60min in the culture medium and imaged with the inverted fluorescence microscope. After 9-10 days in vitro (DIV) healthy slices flattened to approximately 100μm and were selected for further experiments.

### Cerebellar and microglial culture treatment

The cerebellar slices were treated either 3 times 24 hours apart with combinations of recombinant TNF (recombinant rat TNF, Biolegend, San Diego, CA, USA), LTα (recombinant human LT-α, Abcam) and IFNγ (recombinant rat IFNγ, Biolegend) at 50 ng/ml, or twice 24 hours apart at a concentration of 100 ng/ml (glutamate levels were always measured 24hrs after the last dose). The microglial cultures were treated either once or twice with doses of 100 ng/ml or 200 ng/ml of TNF, IFNγ, LT-α, TNF+IFNγ and LT-α+IFNγ (each dose was administered every 24hrs and glutamate levels were always measured 24hrs after the last dose). The cerebellar cultures were also treated with conditioned microglial medium after two 100 ng/ml doses of TNF+IFNγ (in this case, each well contained 800 μl of conditioned medium and 200 μl of nutrient medium). In addition, the cerebellar slices were treated directly with glutamate (L-Glutamic acid, Sigma-Aldrich, Merck) twice every 24hours at a concentration of 75μM and 100μM. A group of slices treated with 100 ng/ml of TNF+IFNγ were also incubated at the second dose with 0.6mM of MK-801 maleate (Fu et al., 2009), which is a non-competitive NMDA antagonist (ab120028, Abcam, UK). The culture medium was replaced entirely when the cytokines or glutamate were added to the primary microglia or to the tissues.

### Immunofluorescence of organotypic cerebellar slice cultures

Immunofluorescence analysis of the cerebellar slices was carried out as previously published (Mendoza et al., 2011). The cultures were fixed with cold 4% PFA (1ml added on top of the filter and 1 ml underneath) for 1 hour at room temperature. After fixation, sections were post-fixed with 20% methanol and washed in 0.1M PBS-0.3% Triton X-100. For further permeabilization, slices were incubated with 0.1M PBS-0.3% Triton X-100 overnight. After permeabilization, slices were blocked with 20% bovine serum albumin (BSA, Sigma, Merck) and PBS-0.3% Triton X-100 for a minimum of 4hr at room temperature. Slices were removed from the insert and incubated overnight with primary antibodies (Table 3) at 4°C in a humid chamber. After primary antibody incubation, sections were thoroughly rinsed in 0.1M PBS, three times 10 min each and incubated with the appropriate secondary antibody conjugated to the appropriate fluorochrome for 4 hrs at room temperature. The tissue was rinsed in 0.1M PBS (three times, 10 min each) and dH_2_O, nuclear counterstained with DAPI (diluted in 1:2000) and mounted on slides with Vectashield Antifade Mountant. Coverslips (24×50mm, VWR, International) were fixed to the slides with clear nail polish.

### Glutamate colorimetric assay

The glutamate concentration in the conditioned medium of primary microglial cultures was assayed with the glutamate colorimetric assay kit from Abcam (no. 83389), according to the manufacturer’s instructions. The samples were incubated at 37°C for 30 min and the absorbances read at 450nm using a SPARK multimode reader (Tecan). The glutamate concentration was extrapolated from the standard curve regression equation and the absorbances corrected for background values. Results were expressed in μM and represented by the mean +/- standard error of the mean (SEM) of 3-4 duplicates per condition.

### Data analysis

All the data was processed in R (R Project for Statistical Computing) and plotted with the package ggplot2. For the human tissue data, the rat tissue data and the organotypic tissue cultures, the non-parametric Mann-Whitney test was used to compare groups and non-parametric Spearman’s rank correlation test for correlation between variables. For the rat tissue data, the Kruskal-Wallis test was also used to compare across groups. For the microglial culture studies, the non-parametric Friedman rank sum test was used to compared across treatment groups and timings, and Wilcoxon pairwise comparison with a Bonferroni correction was used to compare between groups (*p<0.05, **p< 0.01, ***p<0.001, ****p< 0.0001). Mean+/-SEM per group was calculated in all the cases and box plots (showing the median, the 25-75% quartiles, the minimum and the maximum), bar graphs or point graphs were plotted.

### Image acquisition and analysis

Images of immunostained sections from the post-mortem human tissue and the rat tissue were obtained with an epifluorescence Olympus BX63 scanning microscope or a SP8 Leica confocal microscope. The cerebellar organotypic cultures were imaged exclusively with the confocal microscope due to their thickness. For the latter, a range of 4-6 z-stacks were taken with an average thickness of 15μm with a step-size of 0.3μm. All images were analysed using Fiji (Image J, NIH, USA) and prepared in Illustrator (Adobe Systems). The quantification of all the cases/samples was performed with the observer blinded to case identification. Regions of interest (ROIs) in human MS tissue were defined as NAWM regions at least 4mm away from a focal demyelinating lesion. MOG immunofluorescence images were taken at 4x to get an overall scan area of the whole block to select NAWM regions. In the post-mortem human tissue, 10 images of HLA-DR+ staining from NAWM regions were taken at 20x. Microglial activation was analysed by quantifying the area occupied by HLA-DR+ microglial labelling microglia by thresholding the images to a specific intensity per image acquired and dividing them by the total area of each image. In the rat tissue, the number of IBA1+ microglia and GFAP+ astrocytes were counted in images taken at 60x magnification.

PNJ disruption was analysed by measuring the length of the axonal paranodal protein Caspr1 immunofluorescence staining in the post-mortem human tissue, rat tissue and cerebellar organotypic tissue slices. Caspr1, Pan-Na, K_v_1.2 and Caspr1-SMI32 triple and double immunofluorescent images were captured with a 63x oil immersion objective. Only Caspr1 positive axons that were in focus were measured in 10 ROIs in the human and rat tissue. In the cerebellar organotypic slices the ROIs corresponded to a range of 4-6 z-stacks located in the regions with a high density of Purkinje cell axons. In rat sections and cerebellar organotypic tissue slices, 200 focused Caspr1-stained paranodes were measured in each preparation. In the post-mortem human tissue, a total of 6800 Caspr1 stained paranodes from 34 blocks of 10 pathologically confirmed cases and 3200 Caspr1 stained paranodes from 16 blocks of 10 control cases were analysed. In the rat tissue, 1000 Caspr1-stained paranodes from the LT-α+IFNγ group, 600 from the GFP group, and 600 from the naïve group were analysed. In the cerebellar organotypic tissue slices, 1200 Caspr1-stained paranodes from 6 cerebellar slices treated with 50ng/ml of TNF+IFNγ, 800 from 2 cerebellar slices treated with 100ng/ml of TNF+IFNγ, and 800 from 4 control cerebellar slices, 1000 Caspr1-stained paranodes from 5 cerebellar slices treated with microglial conditioned medium, 800 Caspr1-stained paranodes from 4 cerebellar slices treated with 75μM glutamate and, 400 Caspr1-stained paranodes from 4 cerebellar slices treated with 100μM glutamate and 1000 Caspr1-stained paranodes from 5 cerebellar slices treated with TNF+IFNγ and MK-801.

For quantification of voltage-gated K_v_1.2 and Na_v_ channel dislocation, Caspr1-K_v_1.2 and Caspr1-Nav double fluorescent images were taken with the 60x oil-immersion objective objective for each human and rat sample. Only positive Caspr1-K_v_ 1.2 and Caspr1-Pan Na_v_ axons were studied, in 10 ROIs, with a minimum of 50 axons per tissue block. RGB line intensity profiles of Caspr1, K_v_ 1.2 and Pan Na_v_ were obtained along the length of the stained axon to assess K and Na channel dislocation. An overlapping region between two signals was defined as the axonal area where K_v_ or Na_v_ channels were located in the paranodal axolemma. Using RGB intensity line profiles acquired with Fiji, intensity signals of K_v_1.2 or Na_v_ and Caspr1 were subtracted from one another per axon. If the difference between both signals was smaller than a variable threshold at the same point in distance, an overlapping region was confirmed. Moreover, the total proportion of axons with overlapping regions was measured in MS and non-neurological human tissue, and in the GFP, LTα-IFNγ and naive rat tissue. The means+/-SEM of all line profiles were calculated per case and across groups (the code used to calculate channel dislocation can be found in the Supplementary Data).

### Computational modelling

A double cable core model composed of 21 nodes was built and solved numerically in NEURON (Hines and Carnevale, 2006) using the backward Euler implicit integration method (McIntyre et al., 2002).The equilibrium potential was set to −80 mV, and the simulations were run at a temperature of 37°C. The code used to generate this model can be found in the Supplementary Data. The geometrical and biophysical parameters used in this model are summarised in Figure 7 B and C. Simulations across the seven different diameters (d core = 0.4, 0.6, 0.8, 1, 1.2, 1.4, 2.7[μm]) were initiated and allowed to reach the resting potential for 1 ms before a current of 2 nA was injected in the mid-point of node 1 as a square pulse of 0.1 ms duration. The relationship between the core diameter, fibre diameter and myelin sheath thickness was taken from electron microscopy studies of the macaque brain (Liewald et al, 2014). The number of myelin sheath was calculated from the myelin periodicity value of 0.0156 μm (Agrawal et al., 2009). The node to node length was taken from the linear relationship measured from rat nerve fibres (117.52 + 30.47 * d_fibre_) (Ibrahim et al., 1995). The juxtaparanodal length was extrapolated from the diameter dependent scaling relationship from the ventral root of cats (19.6 + 2.58 *d_fibre_) (Berthold and Rydmark, 1983). The paranodal length was determined from the average value of Caspr1 staining from the non-neurological control tissue, and the internodal length was derived from the subtraction of two paranodes and juxtaparanodes lengths from the node-to-node length across the seven diameters. The nodal length (1μm) was kept constant for simplicity reasons. Lastly, the peri-axonal space dimensions of the paranode, juxtaparanode and internode were based on the data measured by following dextran tracers in myelinated fibres of mouse sciatic nerves (Mierzwa et al., 2010). The propagation of the AP along the axon was measured up to 10ms (2000 points plotted/ms), and measurements were recorded at nodes 4 and 16 AP and the AP amplitude, width and conduction velocity were measured. AP amplitude [mV] was described as the voltage difference between the most negative voltage during the hyperpolarisation afterpotential and action potential peak (V_max_). AP width [ms] was defined as the time difference at half the amplitude, and conduction velocity [m/s] as the distance between V_max_ of the spikes at node 4 and 16. We used four types of voltage-gated channels were included: Fast Na_v_ channels, Persistent Na_v_ channels, Slow Persistent K_v_ channels, Fast K_v_ channels and Leakage channels. The following voltage-gated channel characteristics were used in our model:

The maximum nodal Na_v_ channel density was set to 1000 channels /μm^2^ (Shrager, 1989; Waxman and Ritchie, 1993) and the single conductance of a fast Na_v_ channel set to 15 pS (Scholz et al., 1993). Thus, the maximal conductance was 1.5 S/cm^2^. Fast Na_v_ channel gating was based on the gating dynamics from measurements of a human nerve used by Schwarz et al. (1995) at 20 °C. The following gating dynamics were based on these experimental data, and the temperature change was adjusted with a q10=2.2 (Richardson et al., 2000; McIntyre et al., 2002; Coggan et al., 2011).

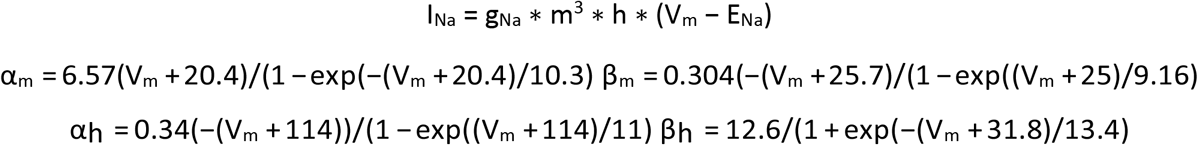

Persistent Na_v_ channel conductance was based on the data from rat ulnar nerves (Bostock and Rothwell, 1997). The estimated density was 6.5 channels/μm^2^ with a conductance of a single channel of 20 pS. Thus, the maximum conductance in this model was set to 0.01 S/cm^2^. This value was taken from previous myelinated axon computational studies (Richardson et al., 2000; McIntyre et al., 2002). The membrane gating dynamics used were the same as previous computational models (Richardson et al., 2000; McIntyre et al., 2002; Volman and Ng, 2013).

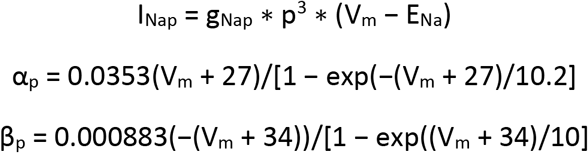

Slow K_v_ channels have an estimated density of 110/μm^2^ (Safronov et al., 1993). From human nerve electrophysiological studies, the single conductance of a channel was quantified to be between 7-10 pS (Scholz et al., 1993; Safronov et al., 1993; Reid et al., 1999). In this model, the single conductance of a slow K_v_ channel was set to 8 pS. Thus, with a density of 110 channels /μm^2^ and the maximum conductance was 0.088 S/cm^2^. The gating dynamics were based on the model of McIntyre et al., 2002.

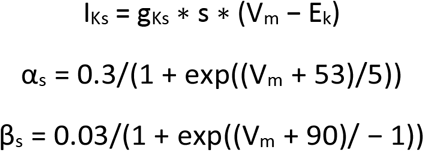

The density of fast K_v_ channels was estimated to be 12 channels/μm^2^ (Safronov et al., 1993) while the single conductance of a channel measured in the rat peripheral nerve was 17pS (Roper and Schwarz, 1989; Safronov et al., 1993). Thus, the maximum conductance in our model for the fast K_v_ channels was set to 0.02 S/cm^2^. These values have been used in previous computational studies (q10=3) (Safronov et al., 1993; McIntyre et al., 2002).

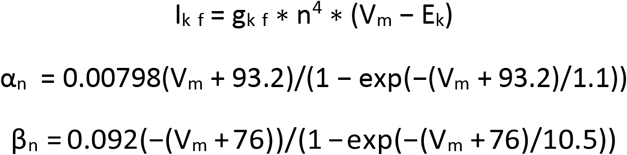

The leak conductance of the node was set to gl=0.007 S/cm^2^ (Bostock and Rothwell, 1997) while the leak conductance of the paranode was gl=0.0005 S/cm^2^, the juxtaparanode and internode conductances were gl = 0.005 S/cm^2^ (Chiu and Schwarz, 1987).

## Acknowledgements

We thank the Centre for Neurotechnology at Imperial College, funded by the UK Engineering and Physical Sciences Research Council, for supporting the PhD studies of PG. The post-mortem human tissue samples were supplied by the UK MS Society Tissue Bank at Imperial College (funding from the MS Society of Great Britain, grant 007/14 to RR). This work was supported by the Multiple Sclerosis Society of Great Britain (grant no. 978/12 to RR and NDM and 037/15 to RR and REJ). NDM and IE were supported by the European Research Council (7th Framework Proof of Concept grant no. 620253). EB was supported by a PhD fellowship from the UK Biological and Biotechnology Sciences Research Council.

---

**Supplementary Figure 1:**
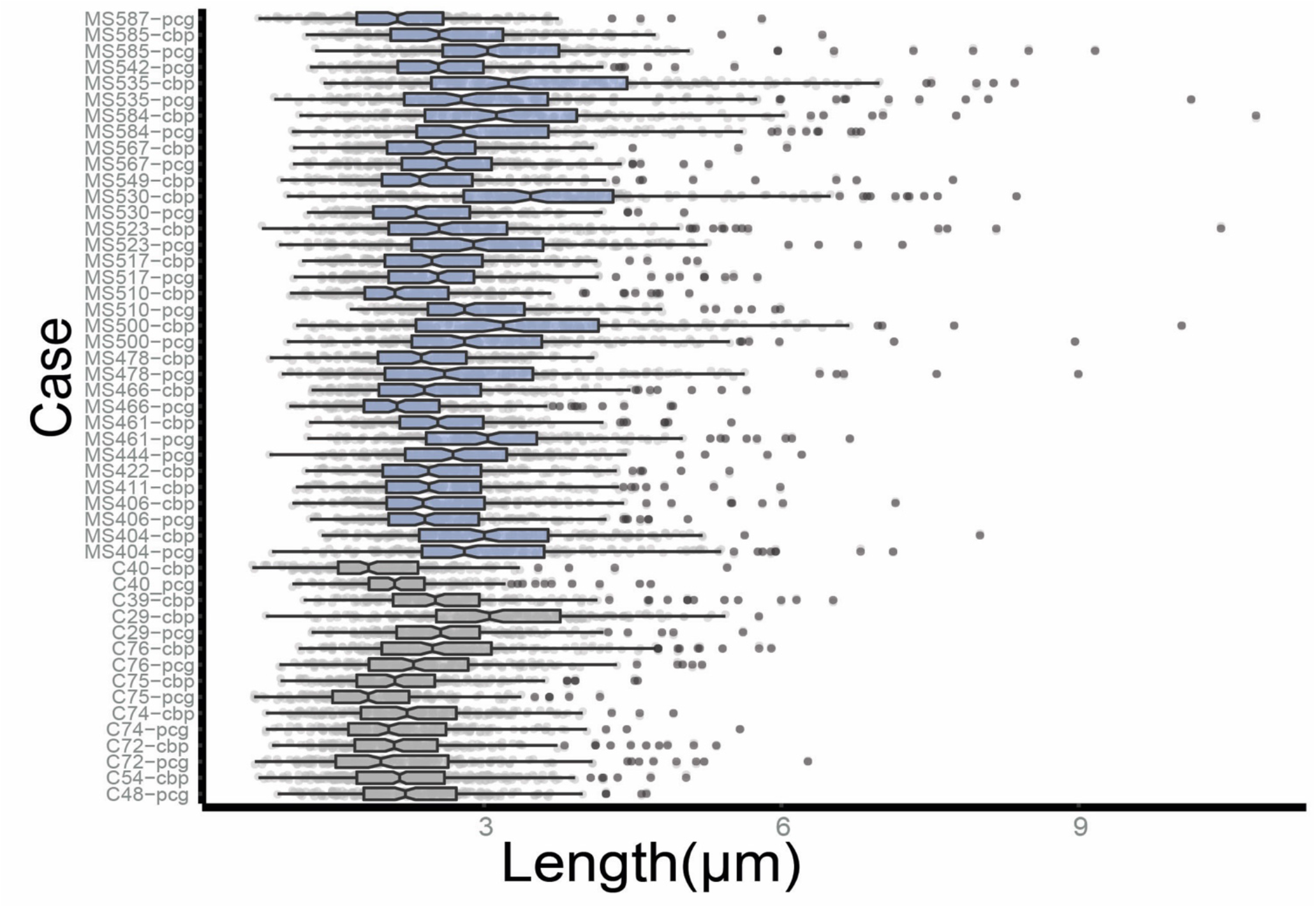
Paranodal Length of MS and non-neurological control blocks. Boxplots representing the distributions of paranodal length from NAWM MS and non-neurological control tissue per case. In the y axis the indexes “cbp” correspond to the cerebral peduncle blocks, while the indexes “pcg” correspond to the pre-central gyrus blocks.

**Supplementary Figure 2:**
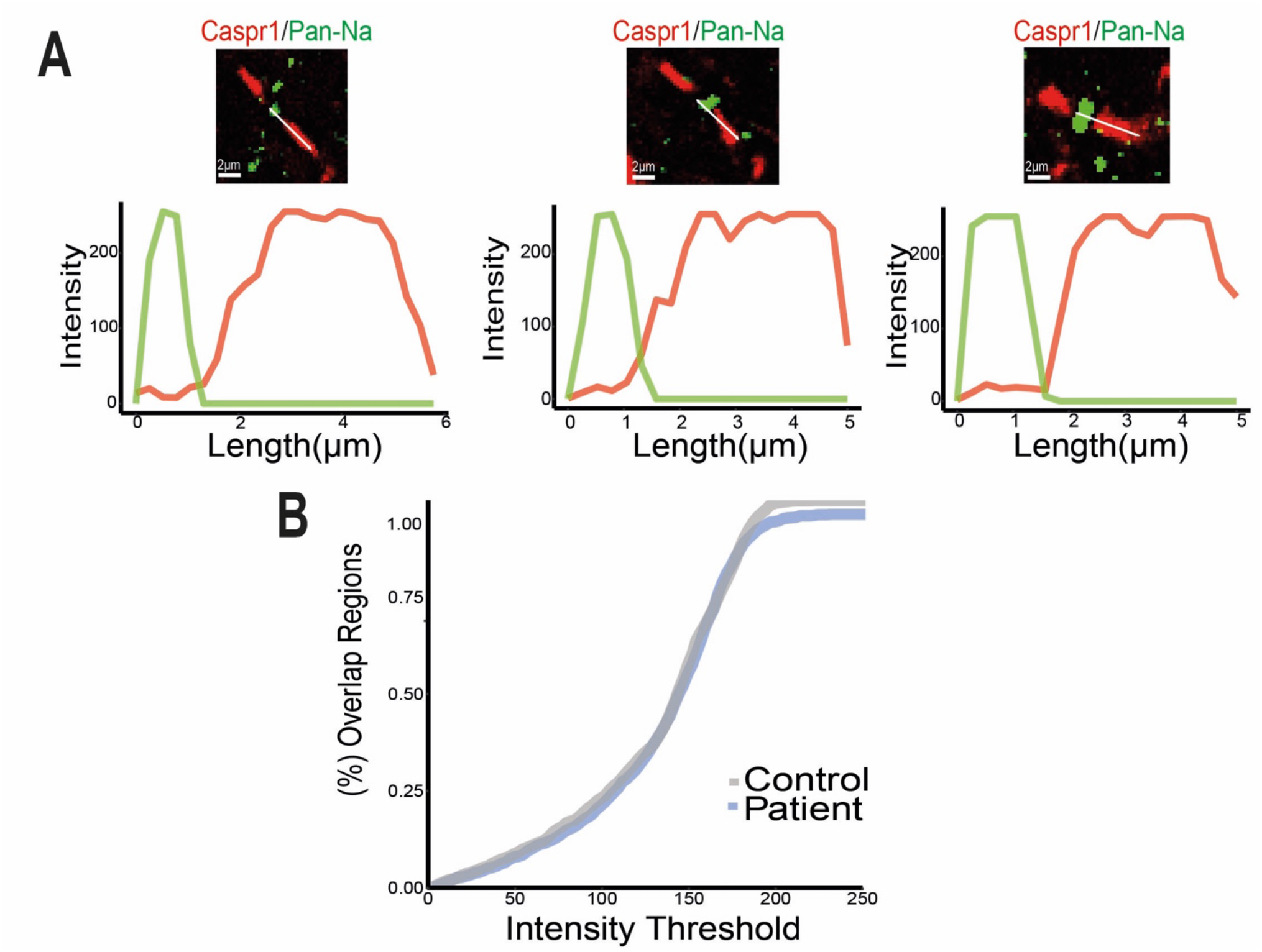
The location of nodal Na_v_ channels was not disrupted in MS NAWM tissue. (A) Confocal images of a double immunofluorescence of Caspr1-stained paranode and nodal voltage-gated Na_v_ channels with the RGB Intensity profile of both immunofluorescence signals across the nodal and paranodal compartments. (B) Caspr1 signal was subtracted from Na_v_, and when the difference between them was smaller than a variable Intensity Threshold, that point was considered an overlapping region. For every threshold calculated, the proportion of overlapping regions was very similar in both groups.

**Supplementary Figure 3:**
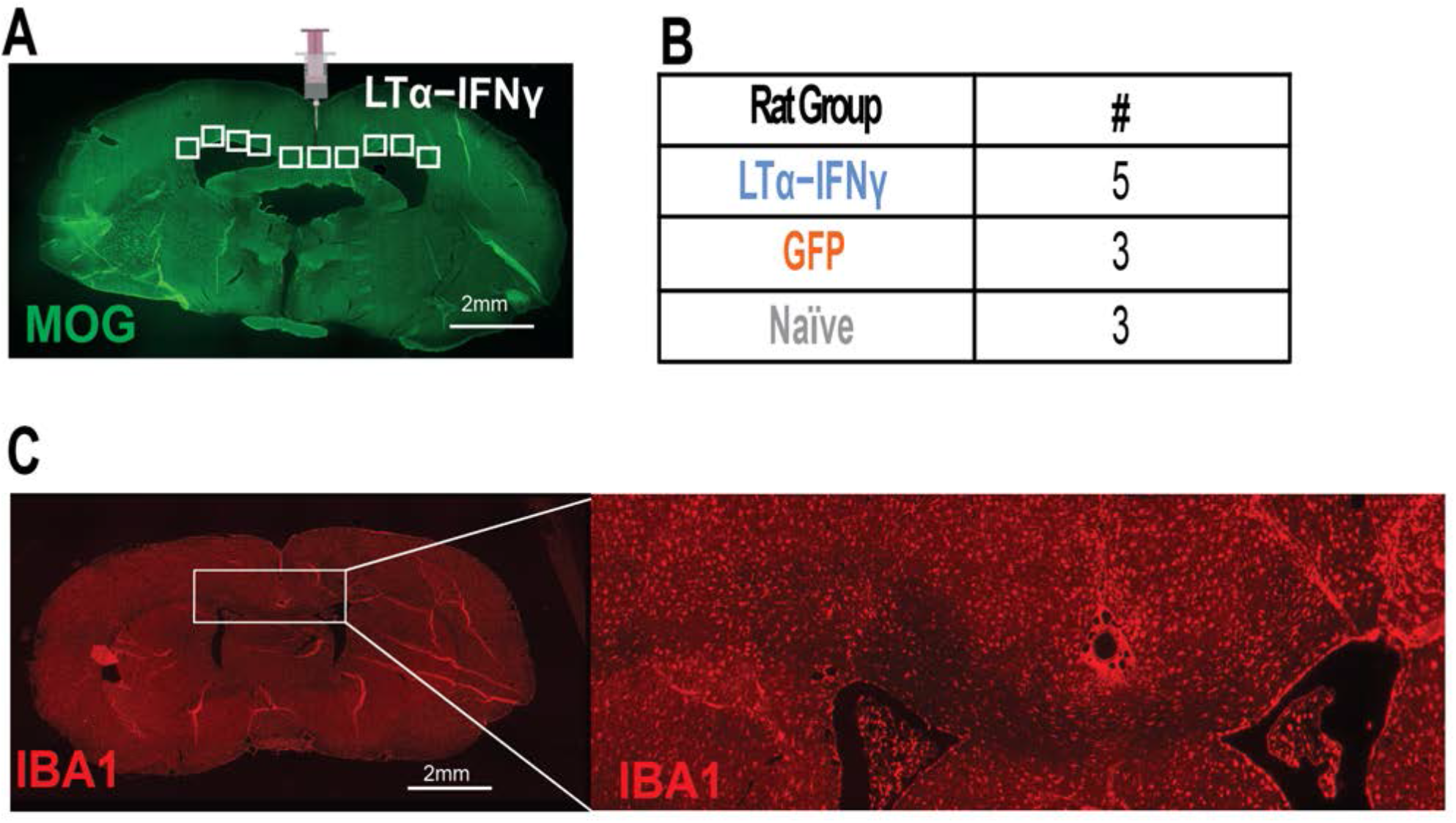
Rat model of meningeal inflammation induced by the chronic exposure to LTα/IFNγ. (A) Immunofluorescent image of a coronal rat section stained with MOG. Lentiviral vectors encoding LT-α and IFN-γ genes were injected into the subarachnoid space in the midline of the brain. The white rectangles are a representative of the 10 selected ROIs at the corpus callosum, cingulum and external capsule. (B) Table of the number of animals used: 5 rats were injected with LTα/IFNγ, 3 rats with GFP and 3 naives. (C) Immunofluorescent image of a coronal rat section stained with IBA1 and treated with LT-α and IFN-γ.

**Supplementary Figure 4:**
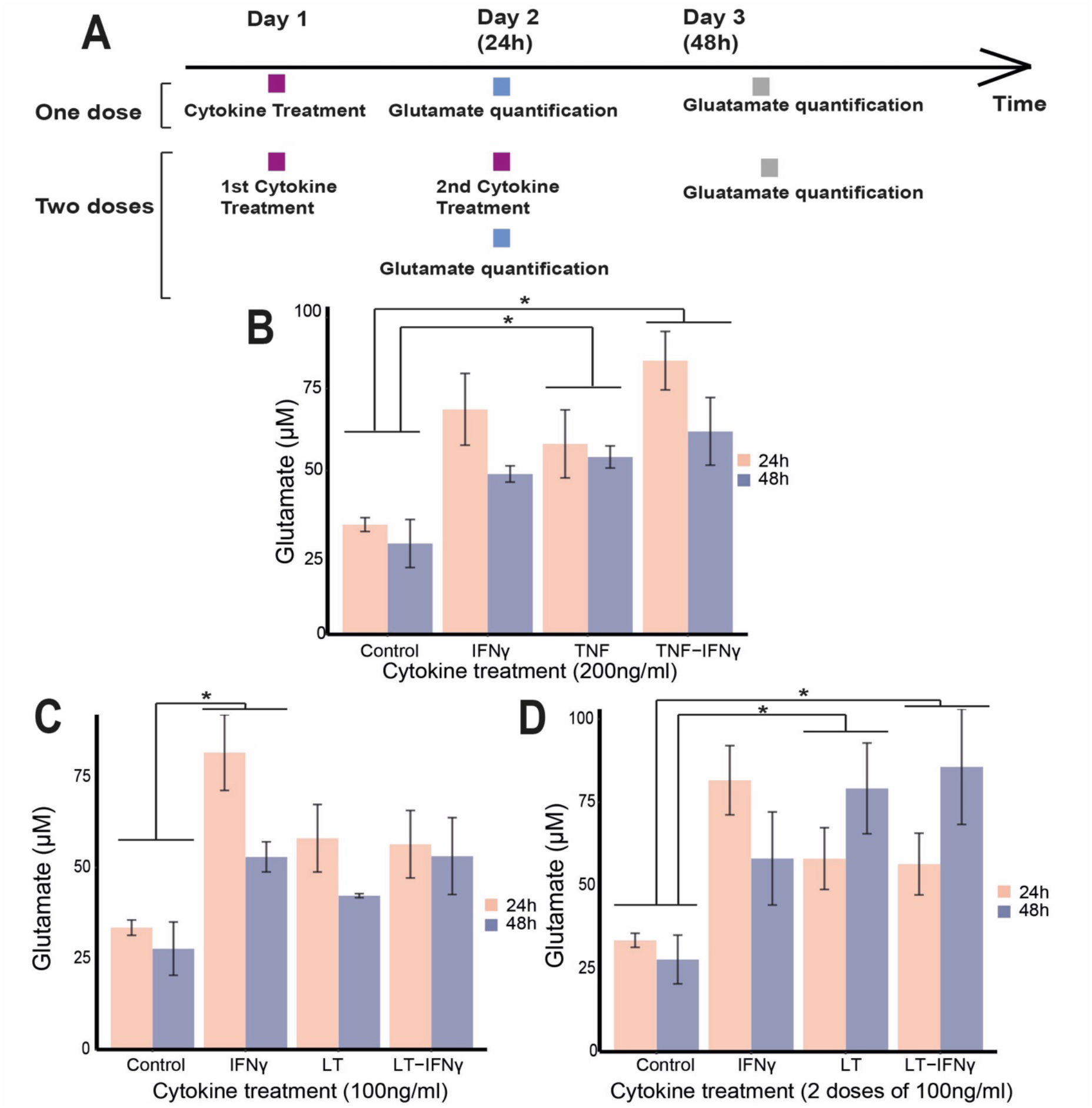
Primary rat microglial cultures images treated with the pro-inflammatory cytokines IFNγ, TNF and LTα. (A) Timeline diagram showing the timings of the experiments. Microglia were either treated with one dose of cytokines or two doses 24 hrs apart. The glutamate in the supernatant was analysed in both cases after 24 hrs and 48 hrs. (B) Mean+/-SEM glutamate levels from replicates showing the statistical difference between controls and the cytokine treatments at different concentrations: 200 ng/ml (n=3 Control, n=3 TNF, n=3 IFNγ, n=3 TNF+IFNγ), (C) 100 ng/ml (n=3 Control, n=3 LTα, n=3 IFNγ, n=3 LTa+IFNγ) and (D) two acute treatments with 100 ng/ml (n=3 Control, n=3 LTα, n=3 IFNγ, n=3 LTα+IFNγ). Non-parametric Friedman test was performed across cytokine groups and timings and post hoc paired-wised Wilcoxon tests to compare groups (* p<0.05, ** p<0.01).

**Supplementary Figure 5:**
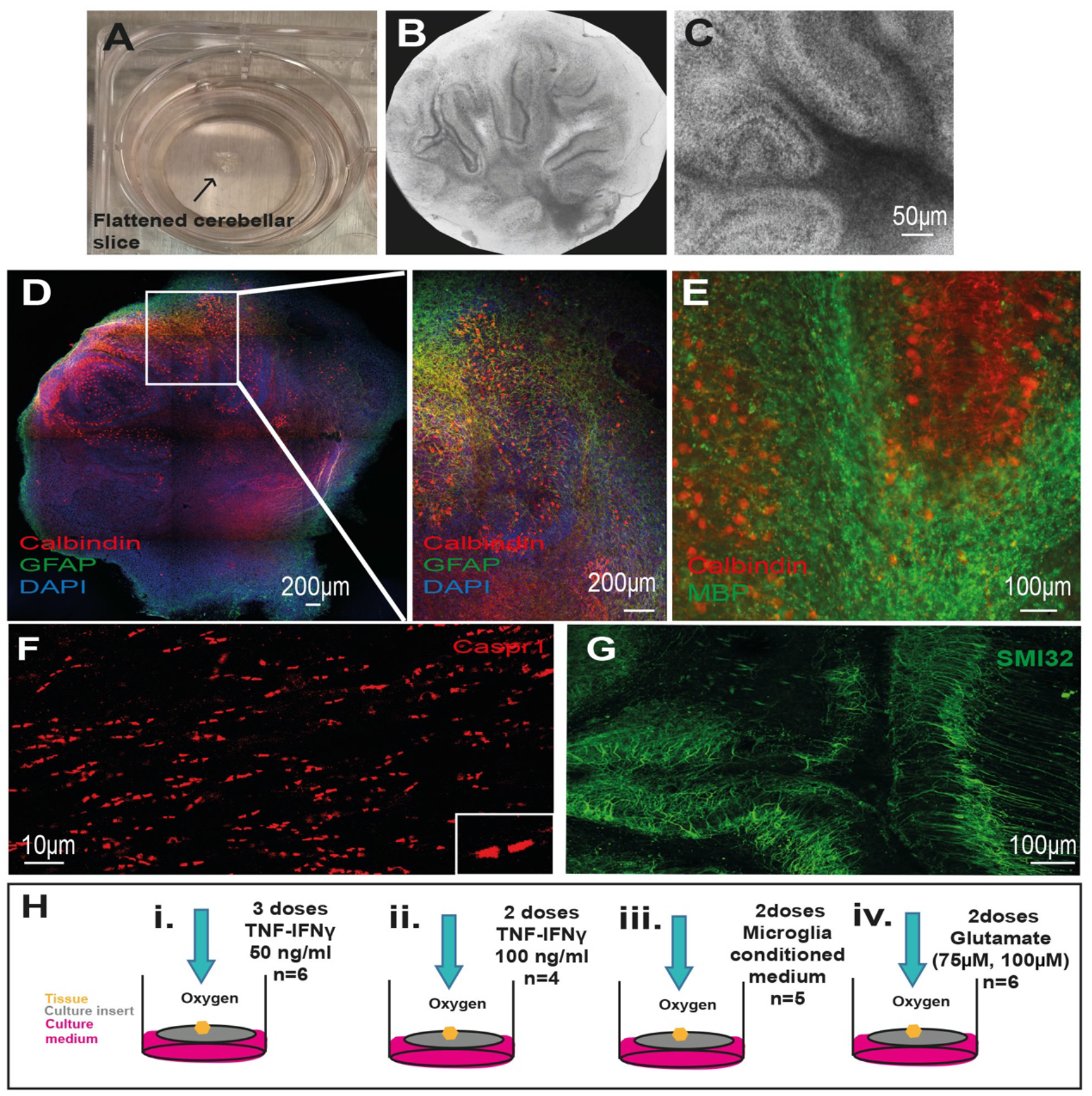
Cerebellar organotypic tissue cultures. (A) Image of a live flattened cerebellar slice. The slices were cut at 400 μm thickness and after 8-10 DIV healthy slices flatten to approximately 100 mm. (B, C) Bright field images of cerebellar slices on culture inserts. (D) Confocal image of a cerebellar slice stained with antibodies against Calbindin+ for Purkinje cells and GFAP+ for astrocytes. (E) Confocal image of a cerebellar slice stained with antibodies to MBP for myelin and Calbindin for Purkinje cells. (F) Confocal image of a cerebellar slice stained with Caspr1 antibodies. (G) Confocal image of a cerebellar slice stained with SMI32 antibodies. (H) Cerebellar slices were treated with the pro-inflammatory cytokines TNF/IFNγ (3 doses of 50ng/ml (n=3), 2 doses of 100ng/ml (n=4)), microglial conditioned medium (2 doses of the medium from microglia treated with 2 acute doses of 100ng/ml of TNF/IFNγ) and glutamate (2doses of 75mM or 100mM).

**Supplementary Figure 6:**
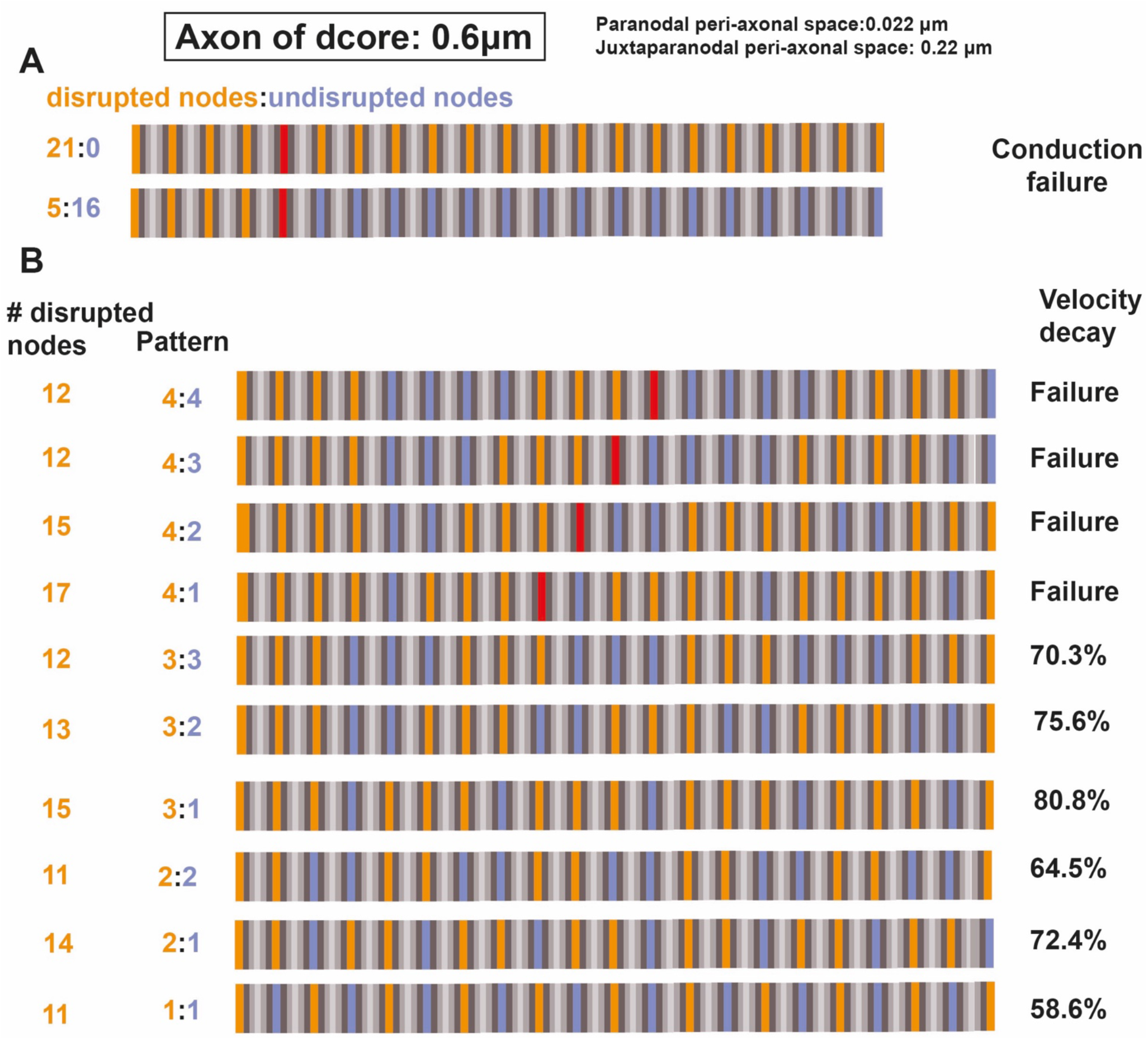
The proportion of disrupted paranodes required for conduction failure. The difference in the proportion of disrupted paranodes within an axon of d_core_ of 0.6 mm can provoke conduction failure and a variable degree of velocity reduction. (A) In axon model of 0.6 mm core diameter, conduction failure occurred when five consecutive nodes were disrupted (orange), and the paranodal and juxtaparanodal peri-axonal space widths were increased up to 0.022 and 0.22 mm, respectively. (B) Velocity decay and conduction failure of this axon model under different patterns of disruption (orange means disrupted node, purple, healthy node and, red denotes conduction failure).

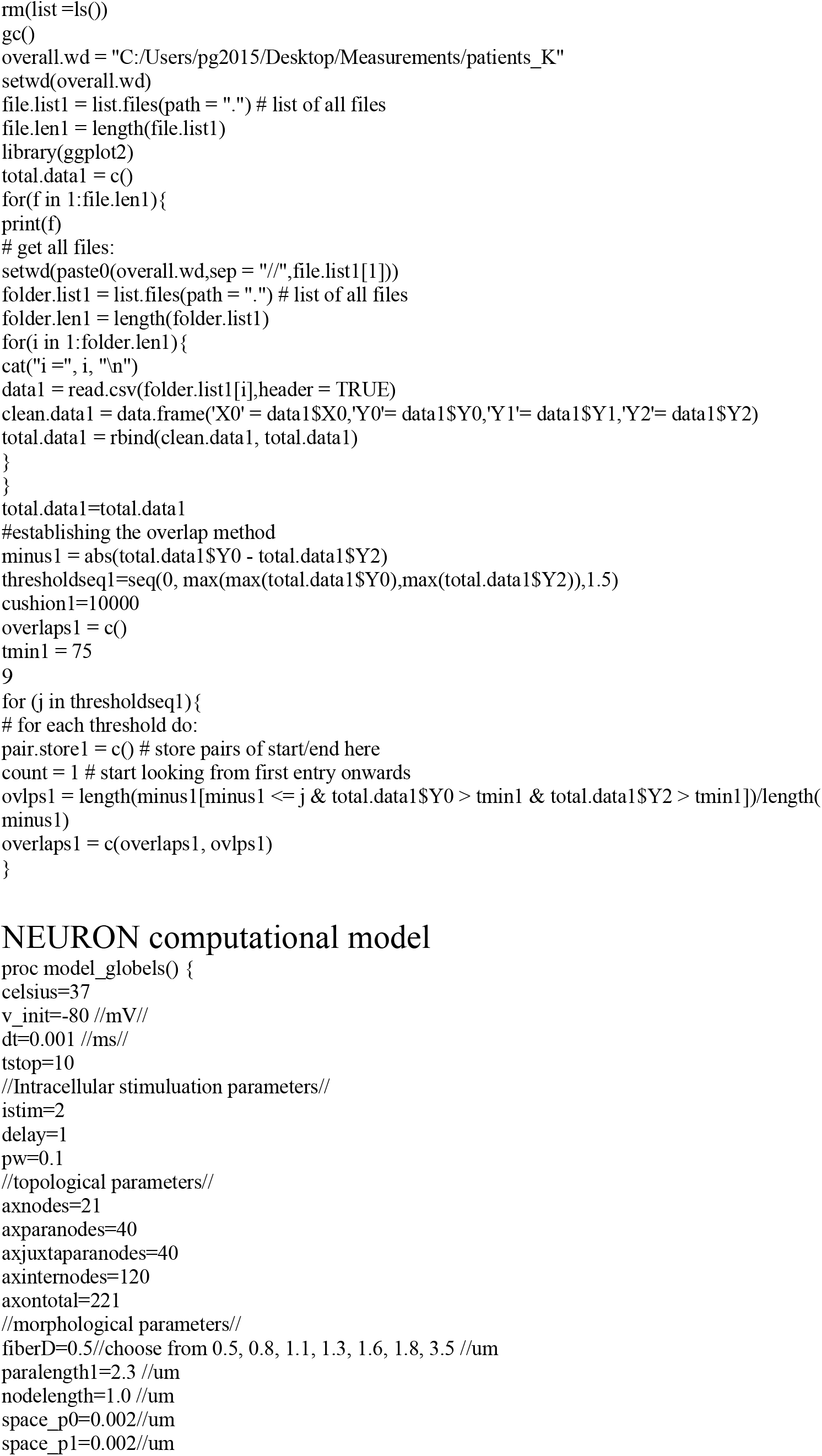
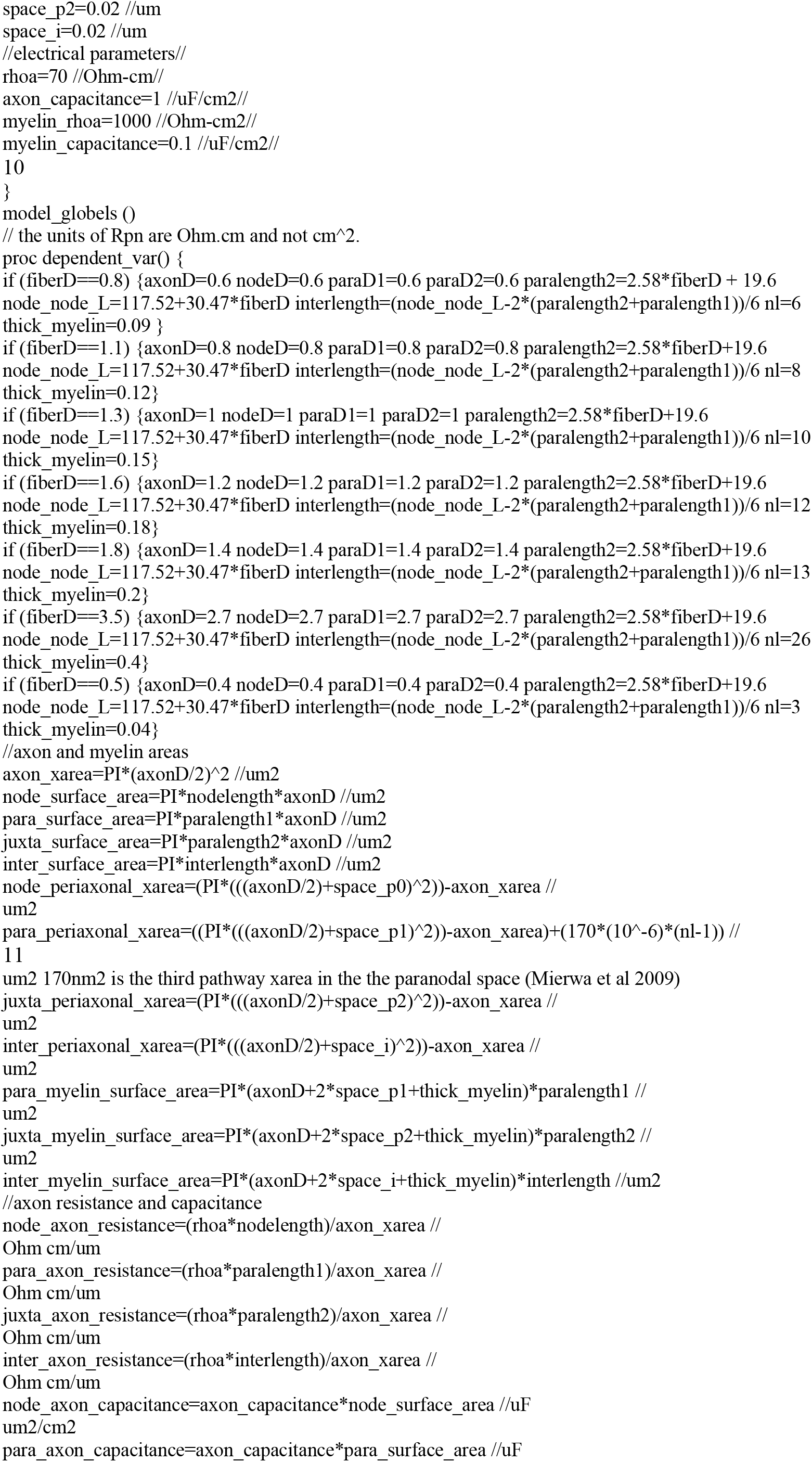
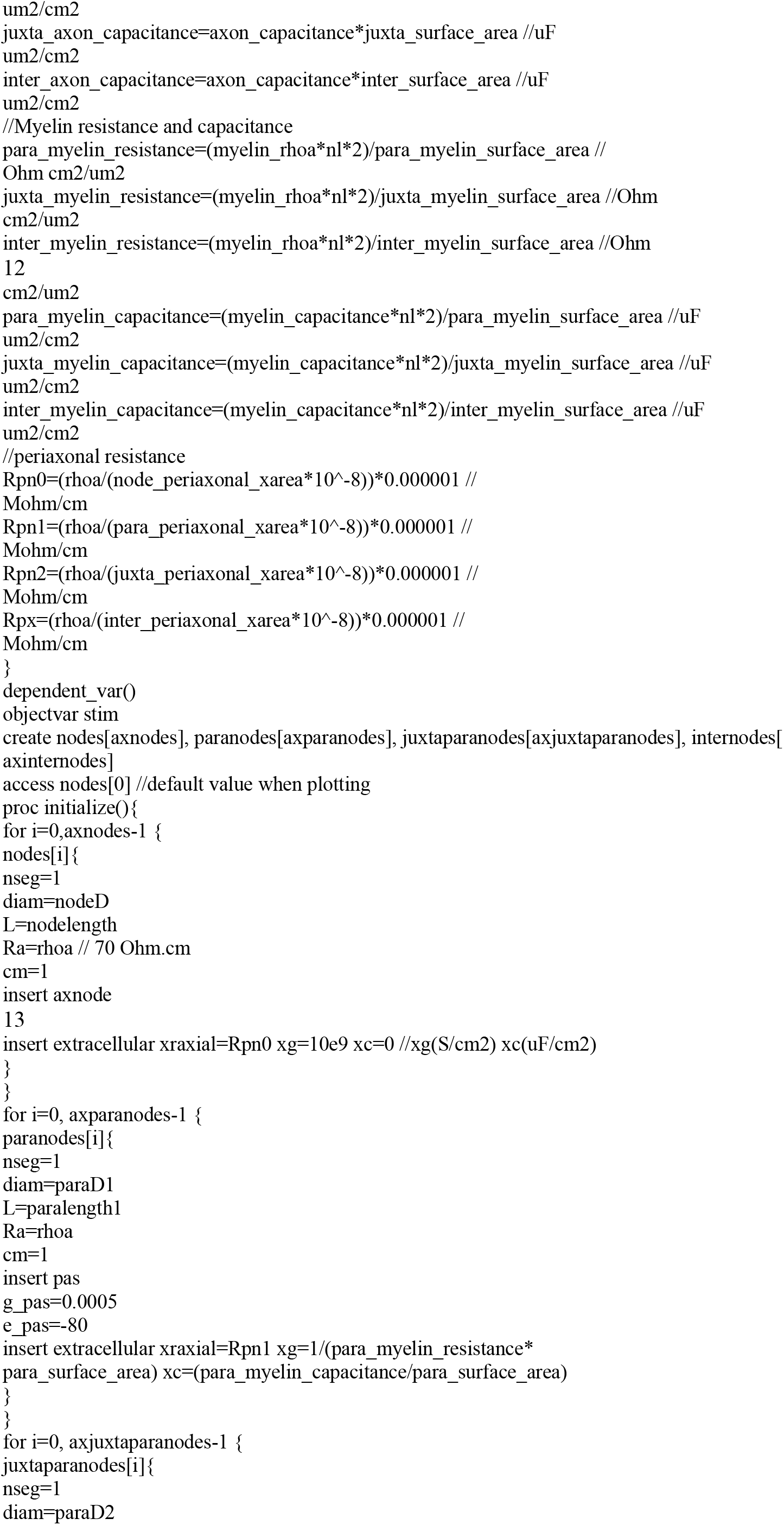
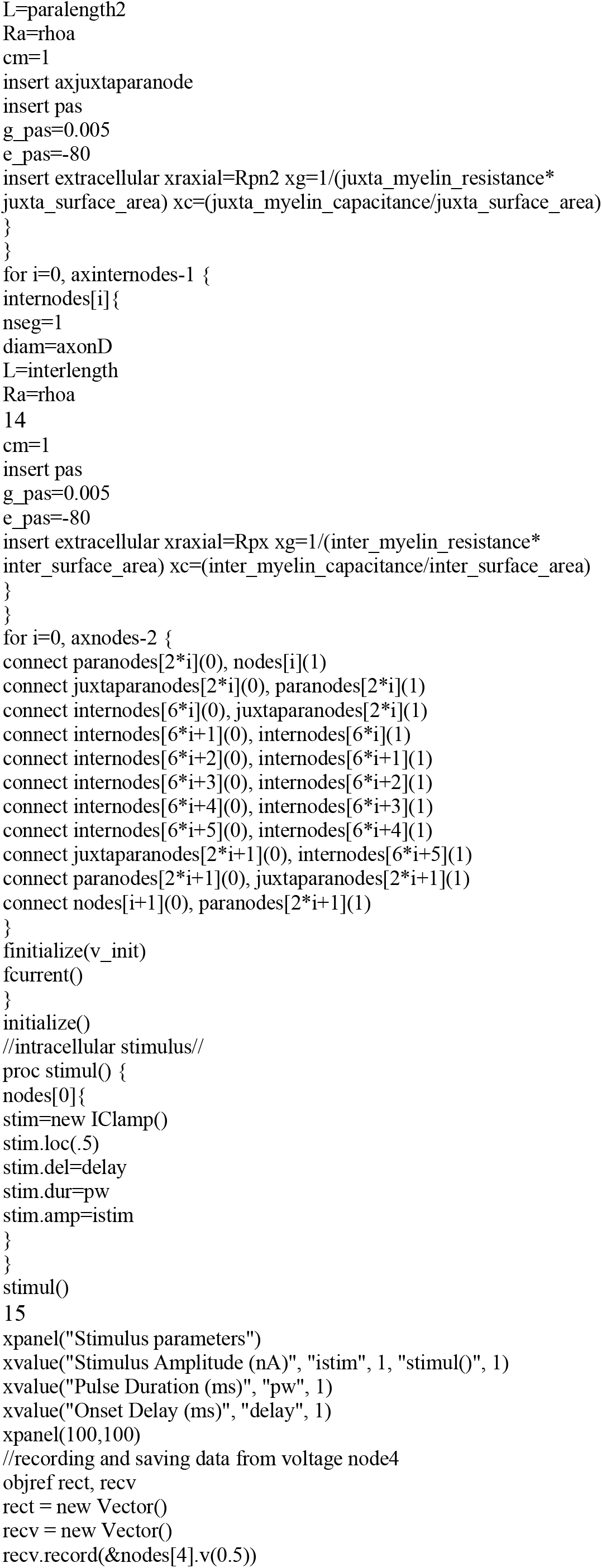
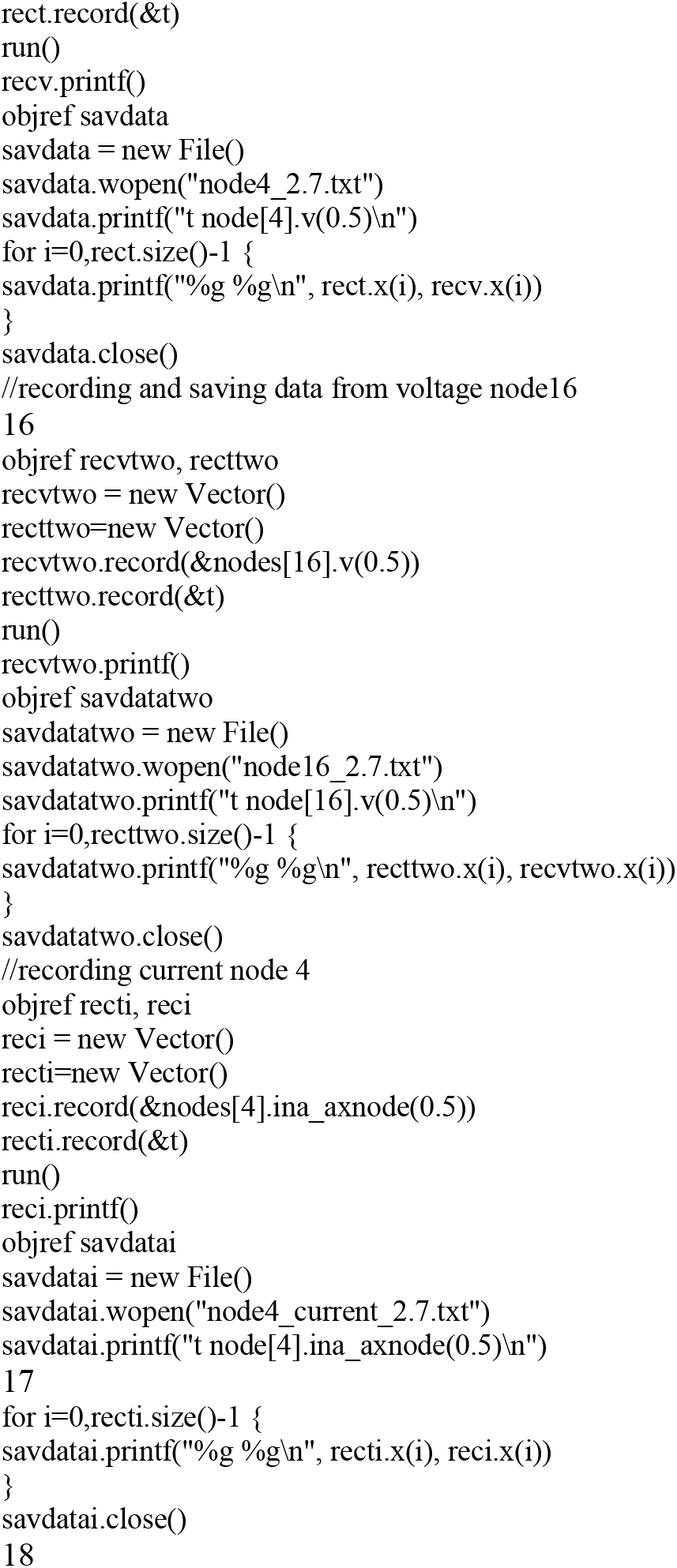
Supplementary Data: Algorithm for the quantification of Caspr1-Kv or Caspr1-Na overlapping signals

